# Systematic Review and Meta-Analysis: Task-based fMRI Studies in Youths with Irritability

**DOI:** 10.1101/2022.01.14.475556

**Authors:** Ka Shu Lee, Cheyanne Hagan, Mina Hughes, Grace Cotter, Eva McAdam Freud, Katharina Kircanski, Ellen Leibenluft, Melissa Brotman, Wan-Ling Tseng

**Affiliations:** Department of Experimental Psychology, University of Oxford, Oxford, United Kingdom; Yale Child Study Center, Yale School of Medicine, New Haven, CT, United States; Division of Psychology and Language Sciences, University College London, London, United Kingdom; Anna Freud National Centre for Children and Families, London, United Kingdom; Emotion and Development Branch, National Institute of Mental Health, National Institutes of Health, Bethesda, Maryland

**Keywords:** dysregulation, fMRI, irritability, meta-analysis, neural activation, systematic review

## Abstract

**Objective:** Childhood irritability, operationalized as disproportionate and frequent temper tantrums and low frustration tolerance relative to peers, is a transdiagnostic symptom across many pediatric disorders. Studies using task-dependent functional magnetic resonance imaging (fMRI) to probe neural dysfunction in irritability have increased. However, an integrated review summarizing the published methods and synthesized fMRI results remains lacking.

**Method:** We conducted a systematic search using irritability terms and task functional neuroimaging in key databases in March 2021, and identified 30 studies for our systematic review. Sample characteristics and fMRI methods were summarized. A subset of 28 studies met the criteria for extracting coordinate-based data for quantitative meta-analysis. Ten activation-likelihood estimations were performed to examine neural convergence across irritability measures and fMRI task domains.

**Results:** Systematic review revealed small sample sizes (median = 58, mean age range = 8–16 years) with heterogeneous sample characteristics, irritability measures, tasks, and analytical procedures. Meta-analyses found no evidence for neural activation convergence of irritability across neurocognitive functions related to emotional reactivity, cognitive control, and reward processing, nor within each domain. Sensitivity analyses partialing out variances driven by heterogeneous tasks, irritability measures, stimulus types, and developmental ages all yielded null findings.

**Conclusion:** The lack of neural convergence suggests a need for common, standardized irritability assessments and more homogeneous fMRI tasks. Thoughtfully designed fMRI studies probing commonly defined neurocognitive functions may be more fruitful to elucidate the neural mechanisms of irritability. Open science practices, data mining in large neuroscience databases, and standardized analytical methods promote meaningful collaboration in irritability research.

## INTRODUCTION

Childhood irritability (hereafter, irritability), an elevated proneness to anger relative to peers,^1, 2^ has received increased attention in child psychiatry in the last decade. Irritability is characterized by frequent, developmentally-inappropriate temper outbursts, low frustration tolerance, and/or irritable and negative mood.^3, 4^ With an estimated community prevalence of 0.12% to 5%,^5^ epidemiological studies have shown that the negative mental health and life outcomes of irritability extend into adulthood,^3^ predicting risks of major affective symptoms and disorders (e.g., anxious and depressed symptoms)^6–8^ and suicidal ideation/attempts.^9^ Although irritability is a hallmark feature of disruptive mood dysregulation disorder (DMDD), it is a transdiagnostic symptom commonly co-occurring with major psychiatric conditions in youths, including attention-deficit/hyperactivity disorder (ADHD), anxiety disorders, major depressive disorder, and autism spectrum disorder (ASD). This highlights the need to study the neural mechanisms of irritability, which may have treatment implications for many pediatric disorders where irritability occurs.

Over the past decade, many attempts have been made at progress probing the neural mechanisms of irritability using functional magnetic resonance imaging (fMRI). Most of these fMRI studies investigated the brain-behavior association between irritability symptoms and task-related blood-oxygenation dependent signals.^1, 2, 10^ The current integrated review focused on three neurocognitive domains in irritability, namely emotional reactivity, reward processing, and cognitive control. Brotman et al.^1^ proposed a translational neuroscience model of irritability that outlined two neural and/or behavioral pathways of irritability— threat processing and reward processing. Evidence for the threat processing pathway showed that when presented with potentially threatening emotional stimuli (e.g., angry and fearful facial expressions), youths with high irritability symptoms and those diagnosed with marked irritability (e.g., DMDD) showed aberrant reactivity in subcortical regions, such as the amygdala, insula, and thalamus, relative to typically developing peers (e.g.,^11–13^). These aberrant neural responses are thought to reflect heightened threat responding in youths with high irritability.^1, 13^ Here, the term emotion reactivity was used given that task fMRI studies in the field commonly compare neural responses to threat or negatively-valenced stimuli versus positive and/or neutral stimuli.

Most evidence for the reward processing pathway was grounded in frustrative nonreward, a negative valence construct in the Research Domain Criteria (RDoC)^14^ matrix. When the omission of expected reward elicits frustration, youths with high irritability showed aberrant neural responses in fronto-striatal regions, such as the prefrontal cortex, cingulate gyri, and caudate, compared to typically developing youths.^15, 16^ Other studies also tested reward processing without the use of rigged reward schedule to evoke frustration, and reported less consistent results in the frontal^17^ and temporal gyri.^18^ Together, aberrant fronto-striatal responses, notably those elicited by frustrative nonreward, are conceptualized as deficits in reward-related processing underlying irritability.^1^

A smaller body of task fMRI studies investigated cognitive control-related functions, probing the top-down regulation and coordination of cognitive processes. These studies have found that youths with high irritability symptoms showed inhibitory deficits, and that irritability symptom severity was associated with aberrant activation in the superior frontal and temporal gyri, inferior frontal gyri, and anterior cingulate cortices during inhibitory control tasks.^19, 20^ According to the exposure-targeted model of irritability,^21^ cognitive control functions facilitate top-down regulation of frustration and outburst behaviors, which are promising targets for intervention.

While these results are promising, there are overlapping as well as distinct regions across these individual fMRI studies targeting different neurocognitive domains. It remains largely unknown if there are convergent neural responses in specific regions that reflect shared neural mechanisms of irritability across threat responding, frustrative nonreward processing, and cognitive control. Also, many past studies had small sample sizes, and variations in research designs (e.g., diagnostic groups, irritability measures, dimensional vs. categorical conceptualization of irritability, experimental paradigms) may limit the generalizability of results and contribute to heterogenous findings across individual studies. Therefore, we conducted a systematic review and meta-analysis to synthesize the irritability fMRI studies published to date to consolidate the current state of knowledge and to identify neural correlates of irritability that are robust to variations in task validity and study designs.

Methodological issues aside, age and sex differences are relatively neglected in the irritability fMRI literature. There is increasing advocacy for attending to developmental differences in pediatric neuroimaging, as developmental stage may moderate socio-affective brain functions.^22^ Fronto-striatal dysfunction following frustrative nonreward was found to be more pronounced in irritable youths in mid-childhood and early adolescence, compared to late adolescence.^16^ However, it remains largely unclear whether the neural correlates of irritability differ as youths transition from one developmental stage to another (e.g., from late childhood to early adolescence when prefrontal circuitries important for mood regulation develop markedly).^23^ Similarly, while research attending to sex differences in irritability symptoms and classification is emerging (e.g.,^24^), irritability studies investigating sex differences in task-dependent neural responses are scarce.

The current integrated review has three major aims. First, we present a systematic review of task fMRI studies focusing on neural activation associated with irritability and related constructs (e.g., reactive aggression, anger) in children and adolescents aged 6 to 18 years, the most common age range sampled in the literature of fMRI research in irritability. By summarizing the sample characteristics and methodological aspects of the studies, we provide an overview of the task fMRI study designs. We also summarize the past studies on age and sex differences in the neural correlates of irritability. Second, we conduct a quantitative meta-analysis based on a subset of qualified task fMRI studies to identify the most robust neural correlates of irritability across neurocognitive domains, i.e., those with high convergence across all individual studies. To provide a more nuanced understanding of the neural mechanisms of irritability, we also examine the extent to which these neural correlates converge specifically within each of the neurocognitive domains examined, i.e., emotion reactivity, reward processing, and cognitive control. Third, we conduct sensitivity analyses to identify potential sources of non-convergence by systematically removing variances due to study heterogeneity (e.g., irritability measurements, dimensional vs. categorical conceptualization of irritability, age differences). We discuss the synthesized results in the context of existing neuroscience-informed models of irritability,^1, 21^ and provide recommendations for future neuroimaging studies on irritability.

## METHOD

### Identification of Task fMRI Studies

A systematic search was conducted based on the PRISMA guidelines^25^ to identify potential task fMRI studies for the purpose of this review and meta-analysis. Importantly, we conceptualized irritability using a transdiagnostic approach, imposing no restrictions on the diagnostic categories of the samples recruited and irritability measures used in the task fMRI studies. Yet, to capture the irritability phenotype as conceptualized, we focused on constructs with marked or highly associated features of irritability, which included anger, reactive aggression, and mood dysregulation.^1, 3, 4^ Such conceptualization hence gave rise to the following search terms and their derivatives: (((irritability) OR (anger) OR (reactive aggression) OR (dysregulation)) AND ((child*) OR (adolescent*)) AND ((fMRI) OR (functional magnetic resonance imaging))), which were used to search for peer-reviewed task-fMRI journal articles published in English, from January 2000 to March 2021. The systematic search was run in PubMed, PsycINFO, Medline, and Web of Science. To ensure that the search included all the key fMRI studies of interest, the identified list of articles was cross-checked with a recent narrative review on the neural dysfunctions of irritability.^1^ Details of the screening procedures and information regarding the exclusion of articles were outlined in the PRISMA flow chart (Figure 1). After independent screening, in-depth reading of full articles, and consensus meetings with senior authors, a final collection of 30 articles were included in the systematic review (Table 1), 28 of which included whole-brain analyses and thus qualified for the quantitative meta-analysis. The identified studies were published between 2009 to 2021, 20 of which were published after 2015. Upon independent data extraction, three of the 28 studies were further excluded from the main quantitative meta-analysis because significant clusters were found in the ROI analysis only,^19^ no significant clusters were reported for any task interaction effects with irritability independent of age,^26^ and only significant main effects of irritability were found.^27^ This resulted in a final collection of 25 task fMRI studies for the main coordinate-based meta-analysis. A detailed summary of the relevant findings and coordinates extracted from the task fMRI studies can be found in the **Supplementary Materials**. Coordinates were converted to and reported in the Montreal Neurological Institute space using the Yale BioImage Suite. The current review and meta-analysis was registered with the PROSPERO ID: CRD42021253757.

**Figure 1.**
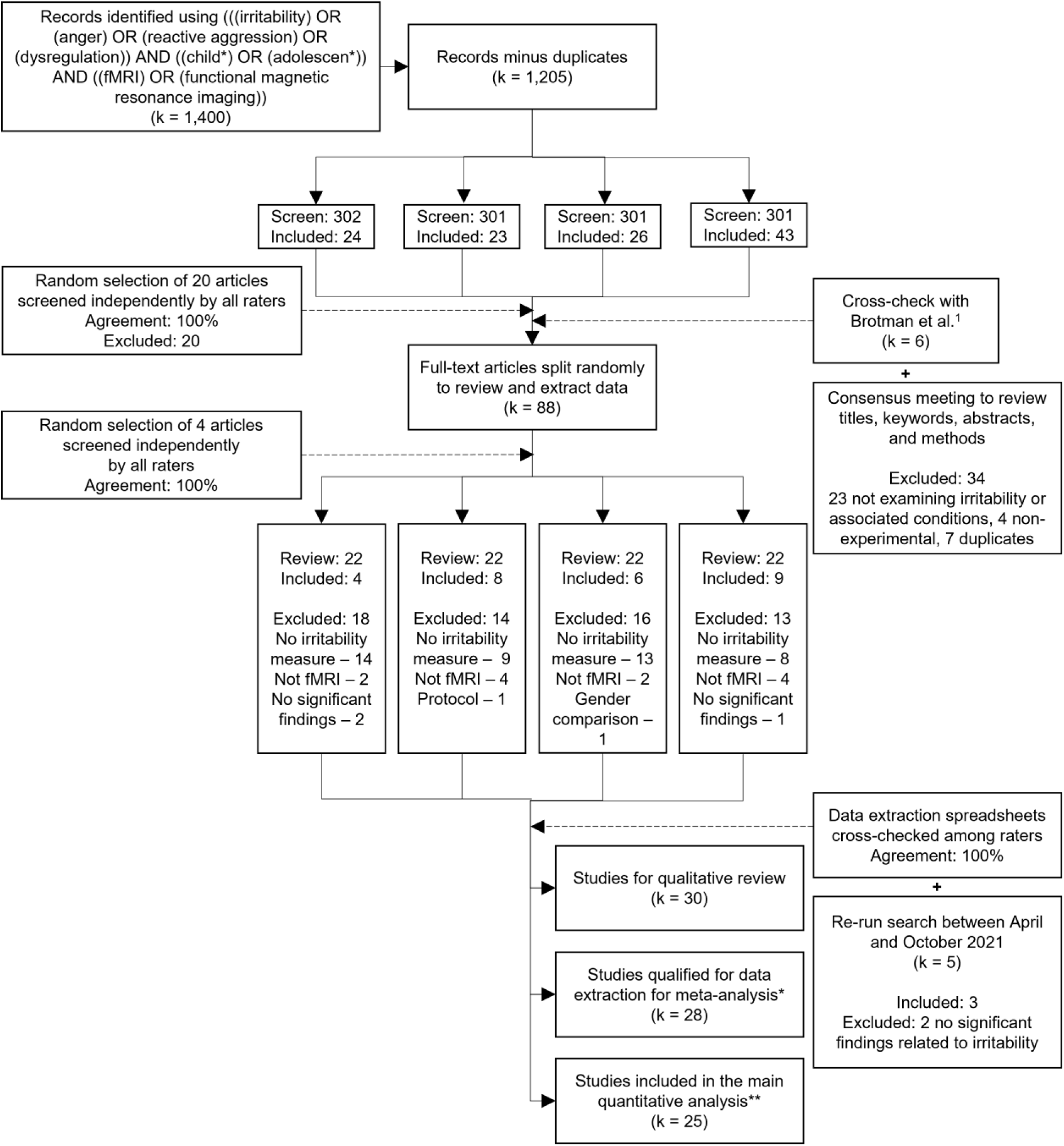
PRISMA flowchart outlining literature search history. *Notes.* *Two studies^28, 29^ were excluded from data extraction because no whole-brain analyses/findings were reported. **Studies reported significant task-dependent neural responses associated with irritability symptoms or irritability-related group differences in the whole-brain analyses across neurocognitive domains (see Methods for further details). Because of the pandemic’s impact on research, the same systematic search was re-run from April 2021 to October 2021 to ensure a comprehensive coverage of studies, and identified three articles that were eligible for the systematic review and meta-analysis.

**Table 1.**
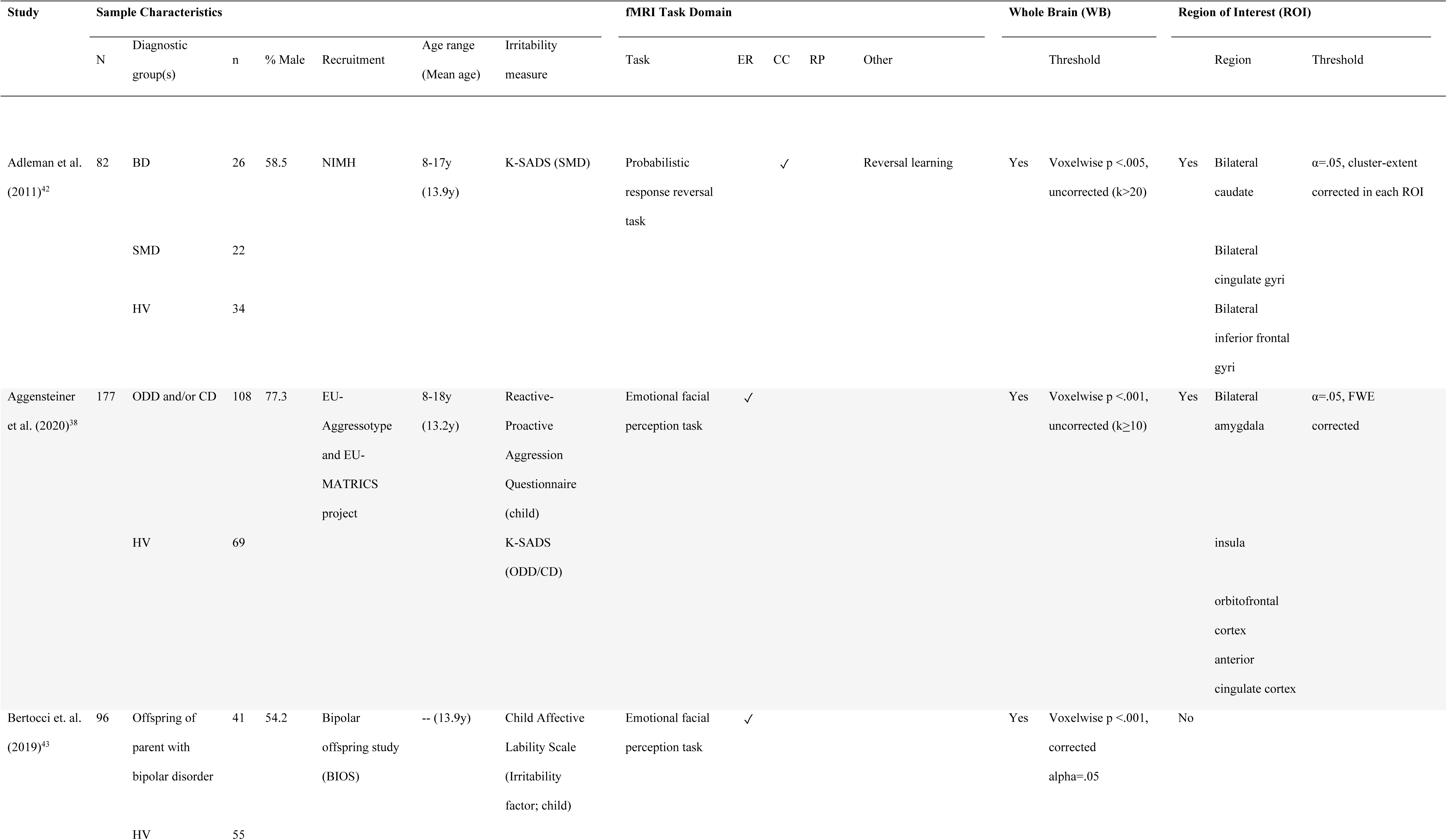
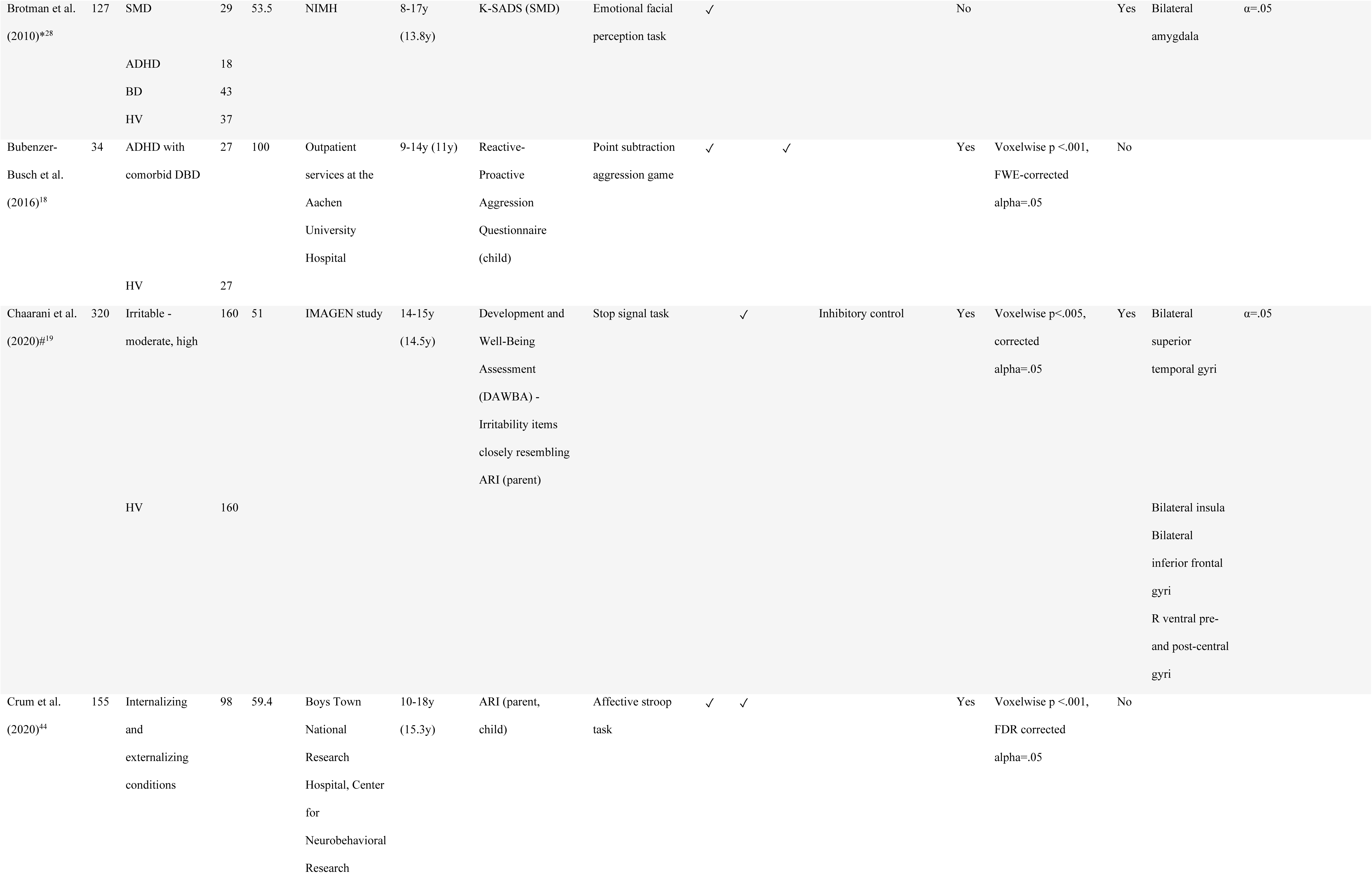
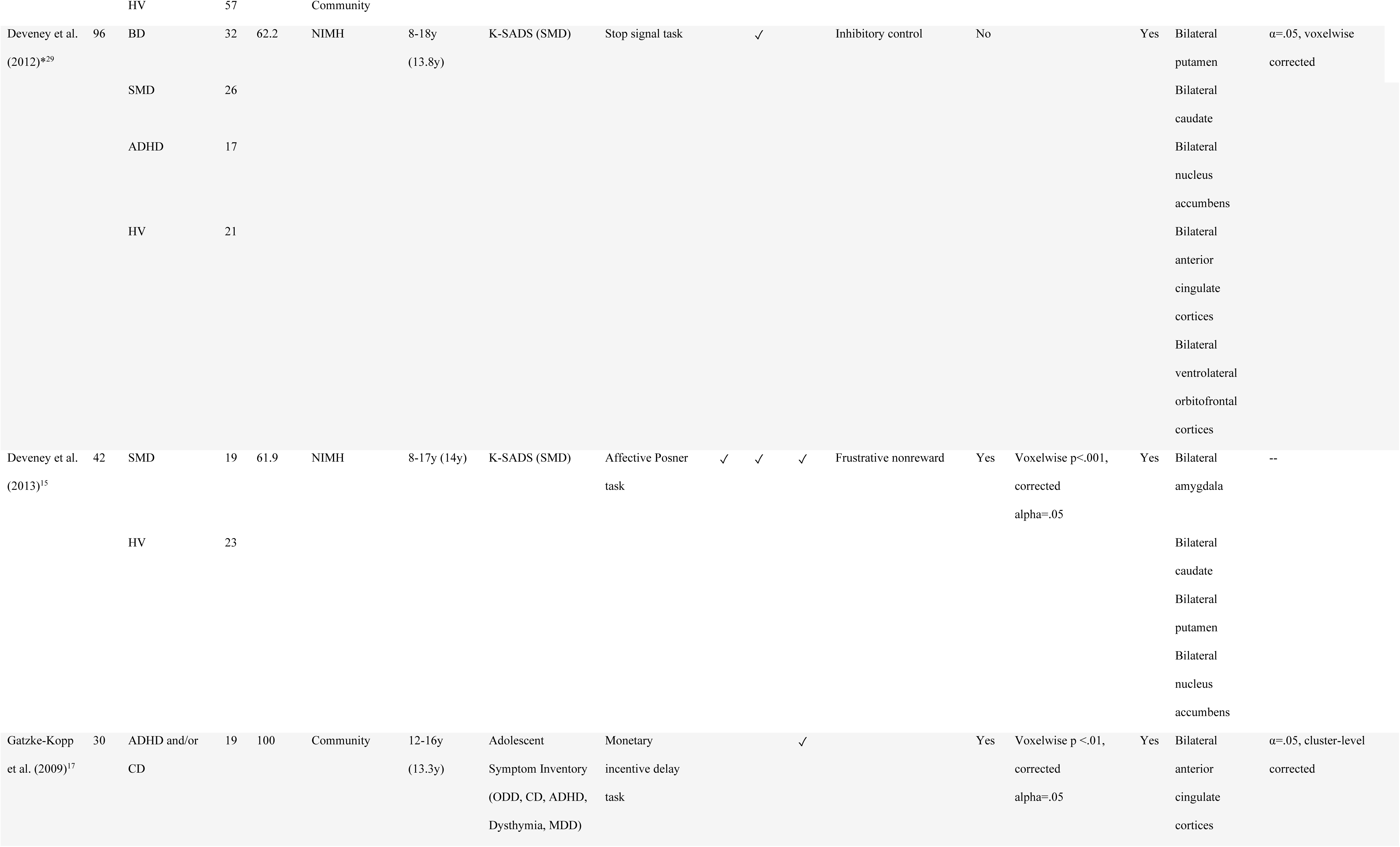
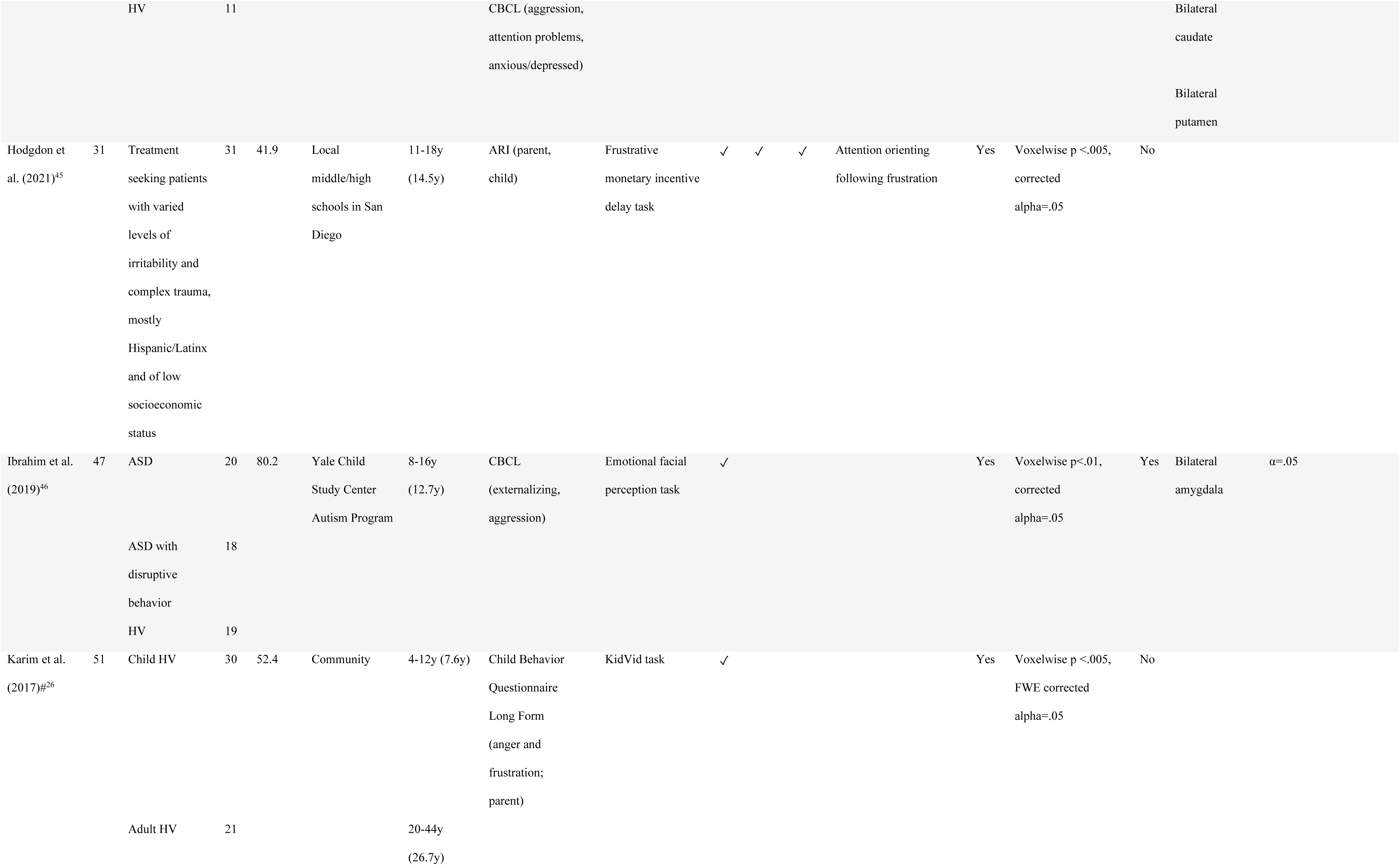
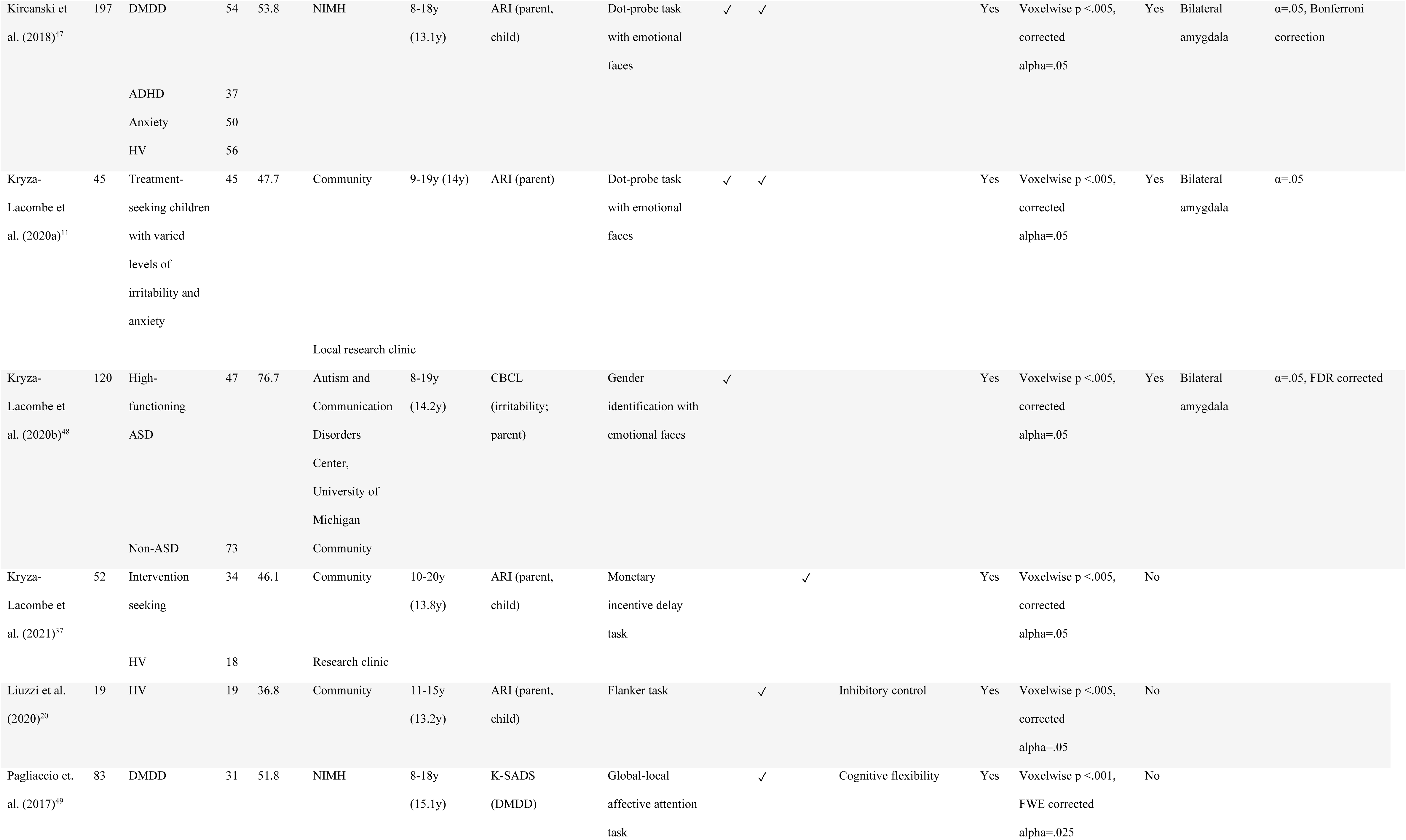
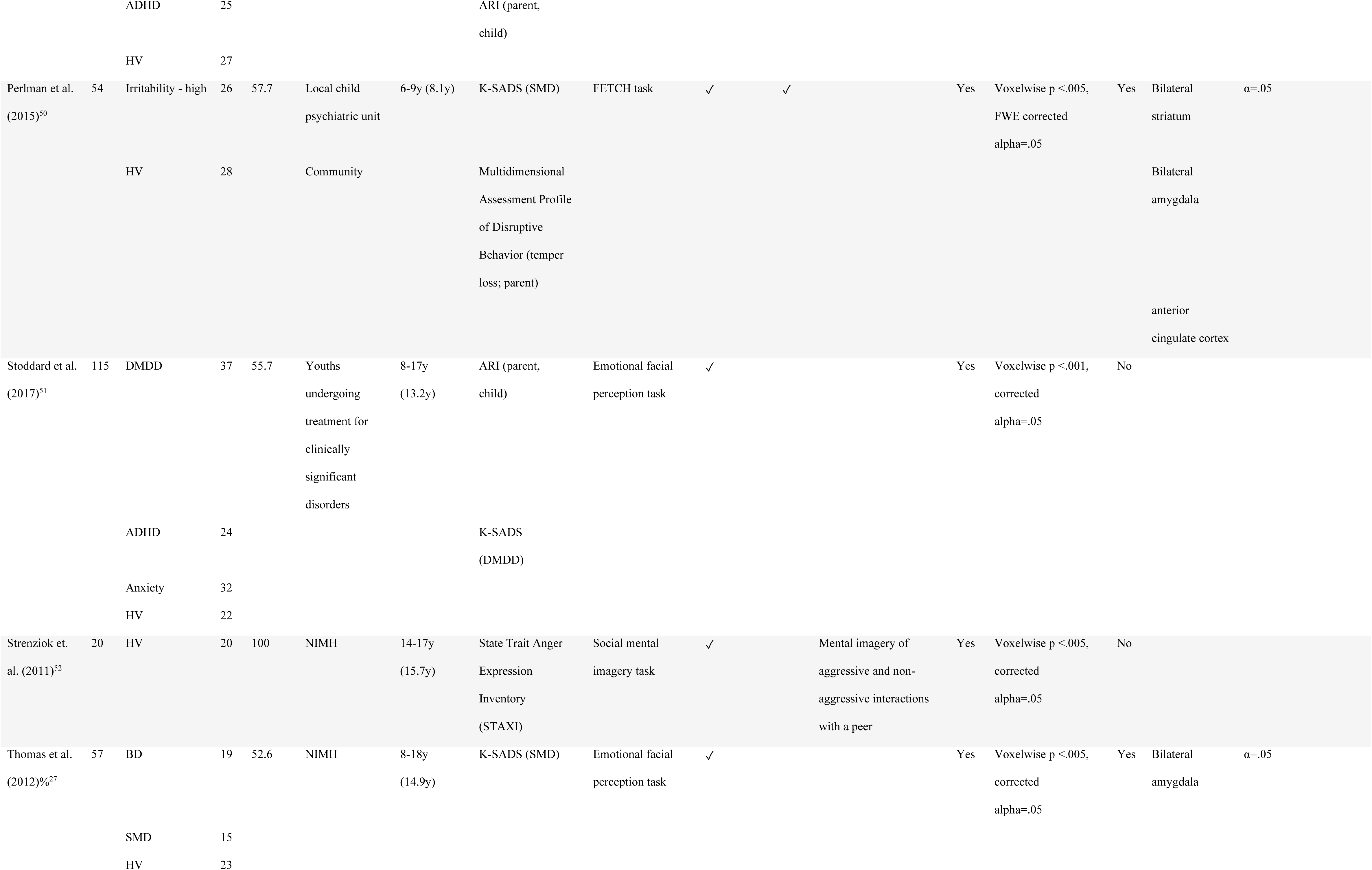
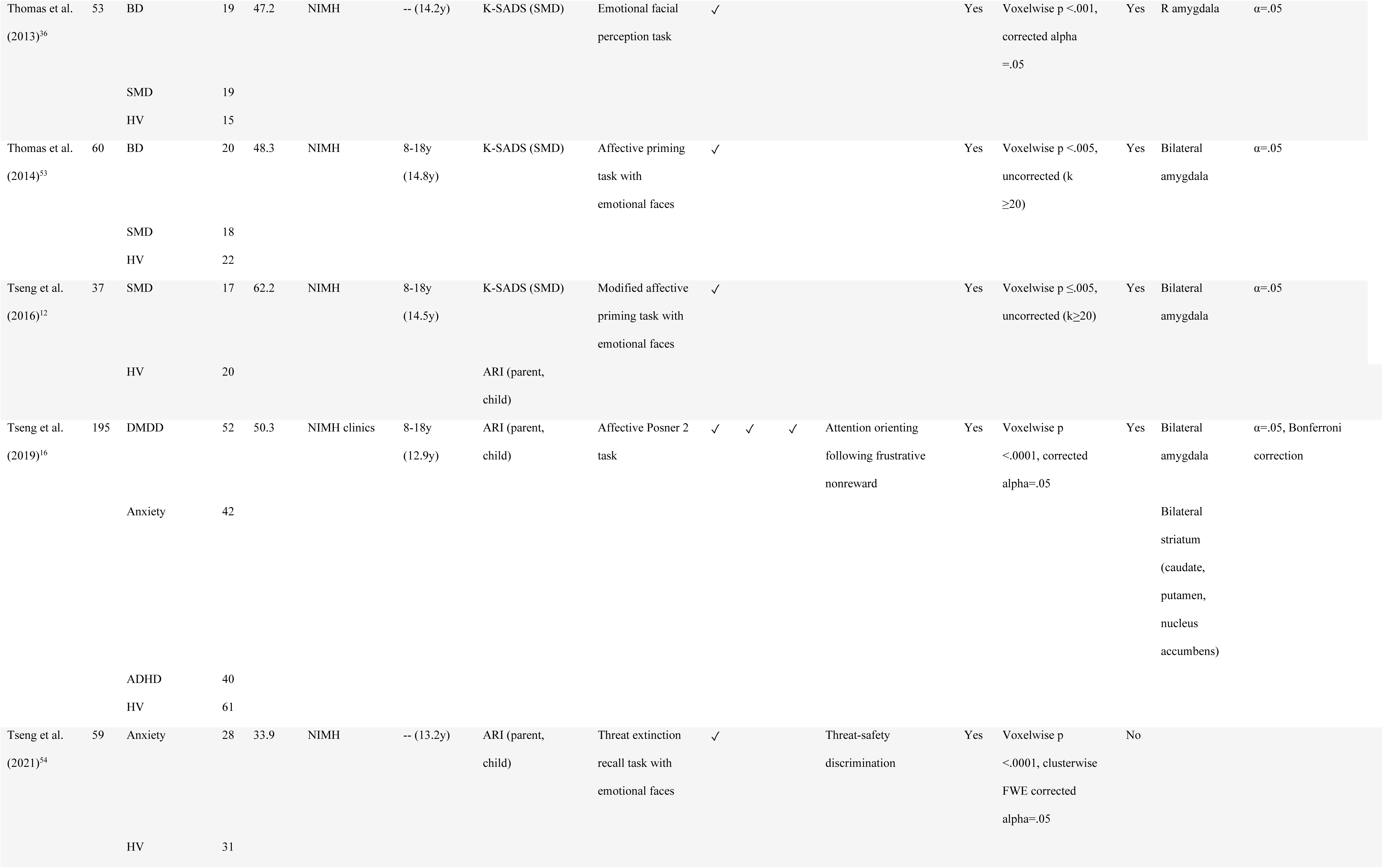
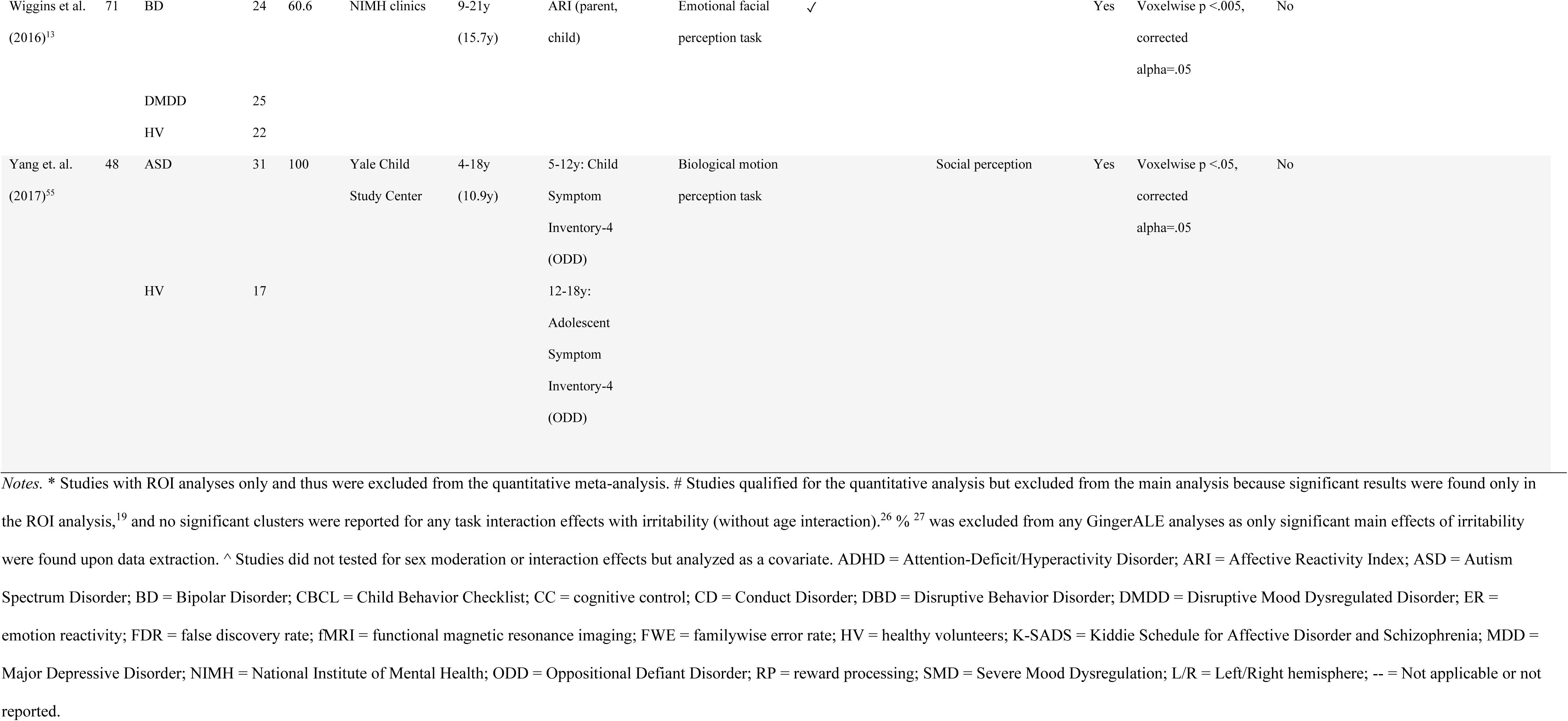
Overview of task fMRI studies in child irritability (*k* = 30)

### Systematic Review

To provide an overview of the task fMRI studies, we first summarized the sample characteristics and key fMRI methodologies reported in the studies. For sample characteristics, we extracted the full and subsample sizes, diagnosis, percentage male, recruitment site, average age and age range, and irritability measure used. For fMRI methodologies, we coded if the studies conducted whole-brain or region-of-interest (ROI) analysis, specific regions of interest (if applicable), fMRI tasks and their neurocognitive domains probed (emotion reactivity, cognitive control, reward processing), and statistical thresholds for conducting those analyses. For emotion reactivity, we categorized studies that employed experimental paradigms that involve the perception of and/or engagement with emotional stimuli. Examples are fMRI tasks that invite participants to view emotional facial expressions or to perform a computer game designed to elicit anger and frustration. For cognitive control, we grouped studies with paradigms that demand top-down executive functions, such as tasks requiring participants to inhibit one’s behavior and orient one’s attention with respect to task demands. For reward processing, we identified fMRI tasks that require participants to engage in reward-driven behaviors, often implemented in a game-like setting along with a reward scheme. We acknowledged that these neurocognitive domains are not completely independent from each other, and it is common that some fMRI tasks might be classified into more than one neurocognitive domain, such as the Affective Posner Task.^15, 16^ Nonetheless, organizing studies by neurocognitive domains allowed for imposing a systematic framework and increasing study availability for the *Subgroup* quantitative meta-analyses, which are insightful for guiding future research. Moreover, we coded if sex differences were examined. Note that two of the 30 studies included in the systematic review did not qualify for subsequent data extraction for the quantitative meta-analysis because whole-brain analyses were not conducted.^28, 29^ Still, a qualitative summary of the sample characteristics and fMRI methodologies of these studies was deemed informative for future recruitment and study design.

### Quantitative Meta-Analysis

Random effects activation likelihood estimation (ALE) was conducted in GingerALE version 3.0.2.^30^ Peak coordinates of the relevant contrasts were extracted from the task fMRI studies and entered to the software, deriving activation likelihood estimates for each voxel. Analyses were conducted where there were adequate numbers of experiments (k=17) as recommended by Eickhoff and colleagues^31^. However, adjustments were made to allow for subgroup analyses of the various neurocognitive dimensions due to study availability (e.g.,^32^). For these subgroup analyses, a minimum of 8–10 studies were required to produce valid results while balancing the need for synthesized fMRI findings with statistical rigor.^32–34^

For our *Main* analysis, a within-group analysis was first conducted using all available task fMRI studies (k=25, 167 foci). Following published guidelines and previous meta-analyses,^31, 32, 35^ statistical significance of the p-value maps was set at a cluster-level inference corrected threshold of *p* <.05, with 1,000 thresholding permutations and an uncorrected *p* <.001. Since including all available contrasts from the identified studies would introduce within-group effects from those that reported alternative analyses of similar contrasts, which could impact the Modeled Activation (MA) values in the software algorithm,^31, 35^ we carefully selected the more interpretable and relevant contrast(s) with respect to the study’s key research interest (e.g., angry vs. neutral faces for facial emotion processing studies;^36^ reward vs. nonreward conditions during reward anticipation, and performance feedback conditions wherever possible for reward processing studies).^37^ For studies that reported more than one relevant contrast with the same control condition (e.g., negative faces vs. shapes and positive faces vs. shapes^38^), the respective coordinates were pooled as one experiment as recommended.^31, 32, 35^ Given that more studies reported significant task-related neural responses when analyzing parent-reported (k=4) than child-reported (k=1) irritability symptoms alone, we prioritized contrasts based on parent report to reduce informant-related variances across individual studies. To gain a deeper insight into the functional significance of the neural aberrations associated with irritability, three *Subgroup* analyses were conducted separately for each neurocognitive domain defined previously. These included *Emotion reactivity* (k=19, 138 foci), *Cognitive control* (k=9, 73 foci), and *Reward processing* (k=7, 52 foci).

Six sensitivity analyses were conducted. First, to supplement the *Main* analysis, we increased the study pool by adding Chaarani and colleagues’ study,^19^ which conducted a whole-brain analysis but only found significant clusters associated with irritability symptoms in the ROI analysis (resulting in a total k=26, 170 foci). Second, we conducted an analysis restricting to only *Emotional reactivity* studies that employed facial emotional processing tasks or involved facial emotion stimuli (k=12, 92 foci), given the relatively large number of such tasks, to reduce task heterogeneity in the *Emotional reactivity* domain. Third, two *Measurement* sensitivity analyses were performed, restricting analyses to studies assessing irritability using the Affective Reactivity Index (ARI^39^) (k=10, 90 foci) and diagnostic modules focused on irritability (i.e., severe mood dysregulation [SMD] and DMDD modules) from the Kiddie Schedule for Affective Disorder and Schizophrenia (K-SADS^40, 41^) (k=8, 59 foci), respectively. These measurement analyses would provide important insights into the potential divergence of neural correlates regarding a dimensional vs. categorical conceptualization of irritability. Relatedly, a *Phenotype* sensitivity analysis was performed by combining the ARI studies with the K-SADS studies (k=17, 137 foci). Finally, a *Developmental* sensitivity analysis was conducted in studies with a mean sample age below 15 years (k=22, 167 foci). We increased the study pool of this sensitivity analysis by adding Karim and colleagues’ work, ^26^ which found significant clusters for an Irritability x Age interaction in a mid-childhood sample (mean age = 7.6 years). Study availability precluded us from conducting an ALE-based subtraction analysis with studies that sampled mid-to late-adolescents (k=4). Of note, although these sensitivity analyses helped reduce heterogeneity, some of these analyses and the *Subgroup* analyses for cognitive control and reward processing had small number of studies and might not capture subtle effects due to limited power. These results should be interpreted with caution.

## RESULTS

### Systematic Review

Sample characteristics

#### Sample size and Age

Across all studies included in the systematic review (k=30), the average sample size was 87 participants (median = 58, *SD* = 66.89, range = 19–320). The number was comparable (mean = 82, median = 55, *SD* = 68.51, range = 19–320) when selecting the most relevant clinical groups with marked irritability symptoms (e.g., DMDD and SMD) for studies that focused on diagnostic group comparisons without dimensional measures. In terms of age, 26 studies recruited pre- and mid-adolescents with mean ages below 15 years (mean = 13.12, median = 13.8, *SD* = 1.89, range = 7.6–14.9), while only 4 studies recruited late-adolescents > age 15 years (mean = 15.45, median = 15.5, *SD* = 0.3, range = 15.1–15.7).

#### Sex proportion

The average sex proportion was 59.8% male (median = 54.2%, *SD=* 17.41), ranging from 33.9–100% (four studies had male participants only).^17, 18, 52, 55^

#### Recruitment

Most study samples were recruited from research facilities with clinical services, such as the National Institute of Mental Health (NIMH; k=14), Yale Child Study Center (k=2), and local psychiatric units (k=5). Four studies sampled treatment-seeking and at-risk youths in the local community.^11, 37, 45^ Two studies assessed irritability symptoms more broadly in healthy community samples.^20, 26^ Three studies constituted part of a large-scale research project (EU-Aggressotype and EU-MATRICS project^38^; Bipolar offspring study^43^; IMAGEN^19^). Based on this summary, it is plausible that several studies might have recruited their samples from the same source (e.g., NIMH) and that there might be overlapping subjects across these studies.

#### Diagnosis

The samples included multiple clinical/research diagnoses: ADHD (n=207, k=8), DMDD (n=199, k=5), BD (n=183, k=7), SMD (n=165, k=8), anxiety (n=152, k=4), and ASD (n=116, k=3), and oppositional defiant disorder (ODD) and/or conduct disorder (CD; n=108, k=1).

#### Irritability measures

Three categories of irritability measures were observed. Ten studies assessed diagnostic categories with marked irritability symptoms using the K-SADS in their main analyses (e.g.,^15, 49, 53^). For dimensional approaches, 10 studies assessed irritability symptoms using the ARI (e.g.,^16^), while 10 other studies used other dimensional measures assessing clinical features associated with irritability symptoms, such as the Child Behavior Checklist (CBCL; e.g.,^48^), Reactive-Proactive Aggression Questionnaire (e.g.,^18, 38^), and Child/Adolescent Symptom Inventory (e.g., ^17, 55^).

### fMRI methods

#### fMRI tasks

A wide array of experimental tasks were employed to probe neural dysfunction pertinent to irritability. Of the 30 studies, 22 studies focused on emotional reactivity, 14 of which involved the perception of and/or engagement with emotional facial stimuli (e.g.,^12, 48^). Seven studies probing reward processing included mostly the Monetary Incentive Delay Task,^17, 37, 45^ the Affective Posner Task,^15, 16^ and other point-based tasks.^18, 50^ Eleven studies probing cognitive control encompassed various subdomains of cognitive control functions in irritability, such as inhibitory control on the Stop Signal Task^19, 29^ and Flanker Task,^20^ reversal learning,^42^ and attention control processes.^49^ Some studies involving emotional reactivity (e.g.,^11, 47^) and reward processing (e.g.,^16^) also probed attention processes (e.g., attention orienting).

#### fMRI analytical thresholds

Heterogenous analytical thresholds were observed across studies. Most analytical thresholds used in the whole-brain analyses were voxelwise corrected (k=18). Other correction methods included those based on familywise error rate (k=5) and false discovery rate (k=1). Four studies reported uncorrected alpha levels. For ROI analyses, similar to whole-brain, most thresholds were not clearly stated (k=10). Other correction methods for ROI included those based on cluster-extent (k=2), Bonferroni correction (k=2), familywise error rate (k=1), false discovery rate (k=1), and voxelwise correction (k=1).

#### Sex differences

Of the 30 studies, only five studies examined sex differences in the task-dependent neural correlates of irritability, and almost all yielded no significant findings (Table S1 in the **Supplementary Materials**), except for two studies that reported a main effect of sex in the left amygdala^27^ and increased activation in several regions important for salience detection during frustrative nonreward processing in younger boys (e.g., insula and pre-/post-central gyri).^16^ Seventeen studies did not report analyzing sex as a covariate nor sex by irritability interaction in their analyses. The eight studies that analyzed sex as a covariate mostly yielded null findings; only one study found sex differences in the salience network during inhibitory control, such as the thalamus and cingulate.^20^

### Meta-Analysis: No Evidence for Convergent Neural Correlates of Irritability

#### Main and subgroup analyses

The *Main* analysis inclusive of 25 task fMRI studies of irritability (167 foci) across all neurocognitive domains revealed no clusters of convergence. Figure 2 visualizes the unthresholded positive z-score map. The three subsequent *Subgroup* analyses focusing on 19 fMRI tasks (138 foci) probing *Emotional reactivity*, nine fMRI tasks (73 foci) probing *Cognitive control*, and seven fMRI tasks (52 foci) probing *Reward processing*, respectively, all revealed no evidence for convergence within domain, suggesting that the null finding in the *Main* analysis was not driven by heterogeneity in tasks across neurocognitive domains.

**Figure 2.**
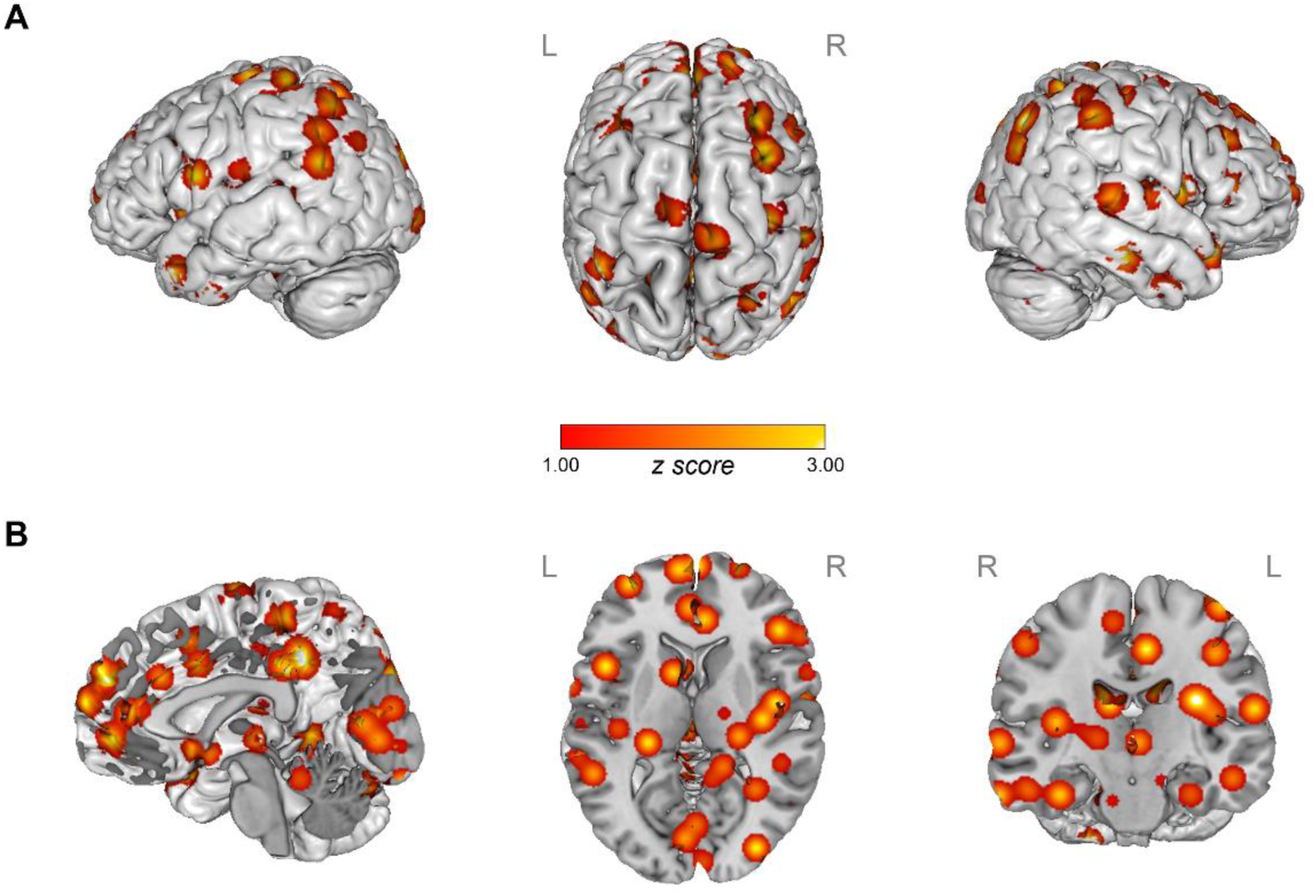
Unthresholded positive z-score map derived from task fMRI studies on irritability (k = 25) *Notes.* Task fMRI studies included in the *Main* quantitative meta-analysis across neurocognitive domains. (A) Cortical regions and (B) Subcortical regions in sagittal, axial, and coronal views (left to right) are presented. No convergent neural correlates of irritability were found across individual studies. L/R = Left/Right hemisphere.

#### Sensitivity analyses

As outlined earlier, six sensitivity analyses were conducted. Given the null findings above, sensitivity analyses may be useful to identify potential sources of non-convergence by systematically removing variances contributed by study heterogeneity. In the first sensitivity analysis adding ROI coordinates from Chaarani and colleagues’ work^19^ to increase the study pool (k=26, 170 foci) and hence power, no convergent clusters were found. Second, restricting the analysis to the emotional face tasks only (k=12, 92 foci) revealed no clusters of convergence. Third, the *Measurement* sensitivity analyses also found no evidence for convergence within the 10 studies (90 foci) that dimensionally indexed irritability with the ARI, and within the 8 studies (59 foci) that analyzed diagnostic categories with marked irritability on the K-SADS. The *Phenotype* sensitivity analysis (k=17, 137 foci) aggregating the ARI studies and the K-SADS studies (which characterized marked irritability using the SMD and DMDD modules) also yielded null results. Finally, the *Developmental* sensitivity analysis on 22 studies (167 foci) with a mean age < 15 years produced no convergent findings.

#### Descriptive ROI findings

Of the 25 studies qualified for the meta-analysis, 15 studies also conducted ROI analyses investigating the association of irritability symptom severity with and/or irritability group differences in task-dependent neural responses in *a priori* defined brain regions. The hypothesized regions were comprised of regions in the salience network underlying the threat processing pathway (e.g., amygdala, insula, and anterior cingulate cortex) and fronto-striatal regions (e.g., inferior frontal gyrus, caudate, nucleus accumbens, and putamen) underlying the reward processing pathway in irritability.^1^ Six of the 15 studies (seven foci) reported significant irritability-related ROI findings. Notably, three out of four studies found youths with high irritability showing increased activation during reward processing^16, 17^ and decreased activation during reversal learning tasks^42^ in the caudate; two out of three studies found increased putamen activation in youths with high irritability during reward processing tasks.^16, 17^ Despite the postulated role of the amygdala in mediating aberrant threat responding in irritability, only two out of 12 studies found increased amygdala responses in youths with high irritability during emotional face tasks.^38, 47^ Figure 3 presents a summary of the ROI findings.

**Figure 3.**
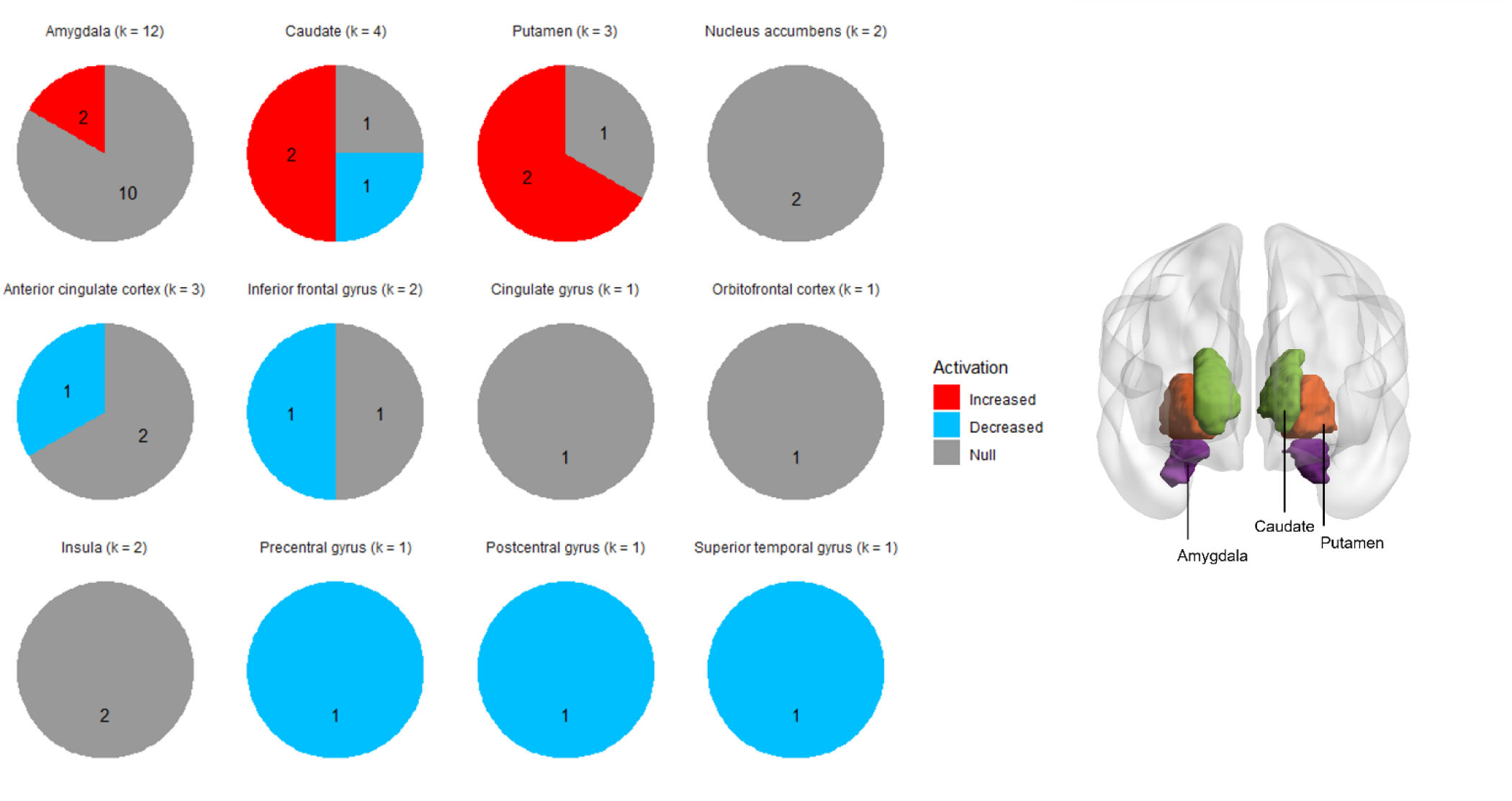
Descriptive summary of ROI findings (k = 15) *Notes.* Pie charts summarize the respective proportions of task fMRI studies that reported an association with irritability symptoms or a related group difference between high versus low irritability groups in each ROI. The brain image depicts the anatomical locations of amygdala, caudate, and putamen, which revealed the greatest number of significant findings across individual studies. ROI = region of interest.

## DISCUSSION

To our knowledge, this is the first integrated and meta-analytic synthesis of task fMRI findings in youths with irritability. We followed the latest recommendations on coordinate-based fMRI meta-analysis^31, 35^ and found no evidence for convergence in the irritability fMRI literature neither in the *Main* analysis across neurocognitive task domains nor in the *Subgroup* analyses for emotion reactivity, reward processing, and cognitive control. Further sensitivity analyses restricting studies by stimulus type, dimensional and categorical irritability measures, irritability phenotype, and developmental ages also revealed no significant convergence across studies. The absence of neural convergence might stem from marked heterogeneity in clinical characteristics, small samples, and variations in fMRI task design, irritability measurements, and statistical procedures, such as thresholding, across individual studies. Moreover, descriptive summary of ROI results suggested altered neural responses during reward tasks in caudate and putamen associated with high irritability, consistent with the striatal reward processing pathway of irritability.^1^

### Heterogeneous Irritability Samples

Although the mean sample size (*N*=87) seemed moderate for neuroimaging research, we noticed considerable variability in the sample sizes; indeed, the *median* sample size (*N*=58) was small across studies. Small sample size not only reduces the power to detect subtle effects, which are common for tasks probing socio-affective processing,^56^ but also may result in inflated estimates, hampering the generalizability of the neuroimaging findings. All of these could contribute to the lack of convergence in the past fMRI studies in irritability.

A myriad of clinical conditions, including DMDD/SMD, ADHD, ASD, ODD, anxiety, and BD, were included in the reviewed studies, which highlights the transdiagnostic feature of irritability. This raises the critical question as to what extent irritability is mediated by similar neural mechanisms across diagnostic categories. We attempted to address this by restricting irritability phenotypes in our sensitivity analysis yet yielded non-convergence. Heterogeneous clinical features and developmental differences in these irritability phenotypes might interact with neurobiological alterations associated with irritability symptoms. Future studies with large samples that are well-powered to examine irritability by diagnosis are necessary to test this possibility.^10^ Moreover, youths with clinical diagnoses are likely to receive psychotropic medications, psychotherapy, and/or have environmental risk factors, such as socioeconomic disadvantages and adverse childhood experiences (e.g.,^45^), which have been shown to alter socio-affective brain functions mediating affective symptoms.^57^ These exogenous factors have not been well-characterized and controlled for in the individual studies, further contributing to the lack of convergence.

Relatedly, studies differ in irritability measures. The most commonly used dimensional measures is the ARI,^39^ while the most commonly used categorical measure is irritability-related modules (i.e., DMDD, ODD, SMD) on the K-SADS. Some other measures included selected items on the CBCL (e.g.,^17, 46, 48^) and Reactive-Proactive Questionnaire (e.g.,^18, 38^). While dimensional measures are more sensitive in capturing individual differences in irritability symptoms and well suited for sensitivity analyses partialing out comorbidity-related variances, categorical approaches allow for identifying the most significant neural correlates in youths with severe forms of irritability warranting clinical attention. Still, there is no gold standard for assessing irritability, and these various measures of irritability differ in measurement validity, reliability, and informant agreement across development.^40, 41^ None of the existing measures are sensitive to low to modest irritability symptoms^40^⎯an issue highly relevant for non-clinical and/or community samples. More justification in the choice of irritability assessments is preferred as irritability-related subscales or items extracted from larger pools, compared to those specifically designed for assessing irritability, might vary psychometrically and relate subtly to different aspects of neural dysregulation.^58, 59^ The measurement heterogeneity and psychometric issues introduce measurement variances to the analysis, which could be another reason for the absence of neural convergence.

Low study availability precluded us from examining age- and sex-related differences in neural convergence. However, we conducted a sensitivity analysis focusing on pre- and early-adolescents only (below age 15) and found no convergent results. Thus, questions remain as to whether age- and sex-related pubertal and hormonal changes might have contributed to the null results, as recent evidence points to an interplay between pubertal hormones and maturation of fronto-limbic circuitries,^60^ overlapping with the threat and reward processing pathways of irritability.^1^ Studies that directly examined sex moderation on irritability-related neural responses are scarce, with only one study found significant sex moderation effects during frustrative nonreward processing.^16^ Together with studies that analyzed sex as a covariate or main effect,^20, 27^ these sex differences primarily emerged in the salience network.

### Heterogeneous fMRI Tasks and Analytical Procedures

Diverse fMRI tasks across multiple neurocognitive domains have been used in past studies in irritability. Although a conceptual framework categorizing studies into emotional reactivity, cognitive control, and reward processing was useful to facilitate systematic analyses of neural convergence, within-domain heterogeneity was still present. This is evident in the emotional reactivity studies reviewed. While most of these studies were fundamentally facial emotion recognition tasks, these paradigms involved varied task demands probing passive and active attentional processes, priming, and control conditions ranging from nose width ratings to gender and shape recognition that potentially involve different psychological processes.

Stimulus variations such as the use of morphed versus non-morphed faces, types of emotions, valence and arousal, and presentation duration might address specific research questions concerning emotion processing in irritability, but that likely further contributes to non-convergence across individual studies given the corresponding impact on the underlying psychological operations and hence associated neural responses. Similarly, a variety of reward tasks were used. Of note, these reward tasks varied in the reward contexts, as some involved the elicitation of frustration via rigged reward (e.g.,^15, 16^) while others occurred in more conventional reward settings (e.g.,^17^). A recent study found task-dependent functional connectivity to be predictive of irritability symptom severity only when frustration was evoked during scan,^61^ highlighting the importance of emotional contexts. There is also inconsistency in operationalizing the temporal dimensions of reward processing in these reward tasks. We strove to reconstruct the full temporal course by carefully pooling study contrasts that reflect the core phases of reward processing (e.g., reward anticipation, reward receipt, and feedback), and yet no significant convergent clusters were found. Studies probing cognitive control are mixed partly because there is no generally-agreed definition of cognitive control dysfunctions in irritability. These subordinate functions range from inhibitory control,^19, 20, 29^ reversal learning,^42^ to attention control processes;^49^ the latter are shared with emotional reactivity and reward processing studies that have attention-related demands (e.g.,^16, 47^). While we do not rule out the possibility that the neural correlates of irritability are indeed very heterogenous due to its transdiagnostic nature and the myriad of neurocognitive functions that are potentially impacted, the heterogeneity in fMRI task designs reflect a lack of consensus in the key neurocognitive constructs of interest and the empirical approaches in probing those neurocognitive processes in irritability research. Study variances related to task heterogeneity are coupled with heterogeneous statistical thresholds in the fMRI analyses. Therefore, the absence of neural convergence is perhaps less surprising.

### Future Directions

Common to many fields of research, bias for publishing novel and significant findings contributes to the use of individualized task designs, flexible preprocessing pipelines, analytical procedures, and thresholding that are unique to individual studies. These research practices often give rise to study findings that are only replicable in well-powered fMRI analyses with sufficiently large samples and representative ranges of irritability symptoms, both of which are difficult to achieve in individual labs. However, this does not necessarily suggest that task fMRI studies on irritability should be replaced with an alternative neuroimaging modality as task fMRI is critical to understanding the functional significance of altered neural functions and their associations with irritability.^1, 2, 10^ In addition to neural activation, task fMRI enables investigation on functional connectivity. Indeed, emerging evidence shows that individual differences in irritability may be reflected in the disrupted integration between and within brain regions and networks.^61, 62^

Several recommendations are noted here for moving irritability fMRI research forward. First, a common battery of agreed-upon irritability phenotype measurements will facilitate comparisons and data pooling sharing across studies and increase sample sizes, potentially improving the convergence of findings. Relatedly, more thorough clinical assessments of co-morbidities would provide the necessary information to clarify irritability-related neural responses that are independent of co-occurring symptoms and heterogeneous features within specific diagnostic groups (e.g., ADHD and ASD). Second, sample characteristics, including information about psychiatric medication, pubertal development, and other environmental risk factors (e.g., chronic stress), are useful to identify exogenous sources of individual variances, enhancing the robustness of fMRI findings. Transparent reporting of potentially overlapping participants, and a wider range of recruitment sites especially in underrepresented populations that are non-white, non-Western and/or at-risk of severe irritability are needed to diversify the study samples. Third, mining population-based neuroimaging datasets, such as the Adolescent Brain Cognitive Development Study (ABCD^63^), provides the opportunity to improve clinical heterogeneity and overcome small sample sizes in individual studies. As measures specifically designed for assessing irritability symptoms are not common in these large-scale studies (e.g., ARI^39^), we advocate for including such irritability measures that are well-validated and reliable in future study protocols. Fourth, fMRI task heterogeneity implies that a better incentive structure is needed to motivate the use of fMRI tasks that validly and reliably probe neurocognitive functions informing the pathophysiology of irritability. This does not mean imposing a stringent framework on fMRI paradigms, as testing novel task designs in individual laboratories are valuable training opportunities for early-career researchers and benefit new hypothesis generation.^64^ Instead, pre-registration of fMRI task designs and analysis plans can promote task homogeneity and standardized processing pipelines across individual studies, while ensuring reasonable between-study variations that address specific research questions. Fifth, open task and data sharing are currently underway in our labs to promote collaborative irritability research. Pediatric neuroimaging in irritable youths can be challenging, especially when frustration tasks and deception are involved. Making mock scan protocols, experimental setups, task instructions, and debriefing procedures openly available may help overcome this challenge. Sixth, the past fMRI studies on irritability were largely conducted at a regional level. Multivariate approaches examining neural coactivation and connectivity patterns across the whole brain may provide a more comprehensive understanding of the neural circuitries and interactions mediating irritability.^61^ Other neuroimaging modalities such as connectivity studies using fractional anisotropy^62^ and functional near-infrared spectroscopy measuring real-time cortical neural responses during interactive tasks^65^ offer novel angles to study neural dysfunctions in irritability. Studies analyzing both task fMRI and task-free resting state data also allow for clarifying task-related neural noises.^62^ Finally, frustration realistically occurs in social and interactive contexts among youths. To enhance ecological validity, future irritability research might investigate neural dysfunctions during frustrative social nonreward, such as social rejection.

### Conclusion

This study is the first systematic review and quantitative synthesis of the task fMRI studies on irritability. We observed vast clinical heterogeneity and methodological variations across studies, potentially contributing to the absence of neural convergence in irritability as shown in the quantitative syntheses across neurocognitive domains and sensitivity syntheses restricting stimulus type, irritability measures, and developmental ages. Nonetheless, when implemented thoughtfully, task fMRI studies are valuable empirical evidence for elucidating the functional neural mechanisms mediating irritability symptoms. The use of large samples, common, standardized measurements of irritability, comprehensive assessments of heterogeneous clinical features, and more homogeneous fMRI tasks probing well-defined neurocognitive domains central to the psychopathology of irritability are key to improving research practice and data quality in the field. Open science and innovative research methods such as multivariate analysis and multimodal neuroimaging provide novel avenues for advancing the current state of knowledge in the neural mechanisms of irritability.

## Acknowledgements

KSL is supported by a doctoral student scholarship awarded by the Hong Kong Jockey Club Charities Trust. KK, EL, and MB are supported by the National Institute of Mental Health (NIMH) Intramural Research Program (ZIAMH002781). WLT is supported by a research grant from the National Institute of Mental Health (R00MH110570) and the Fund to Retain Clinical Scientists from the Yale School of Medicine and the Yale Center for Clinical Investigation. The above funding sources have no involvement in the conduct of the research and/or preparation of the article.

## Availability of data, code, and other materials

Data analysis scripts (GingerALE) are available from the corresponding author upon reasonable request. Data extraction table is provided in the Supplementary Materials for those interested in replicating and/or conducting their own analysis. The protocol and registration information of this review are publicly available under the PROSPERO ID: CRD42021253757. All PRISMA materials (flowchart, checklists, and risk of bias assessment form) will be made available online via Open Science Framework upon acceptance.

