## Supplemental Table S1 for "Systematic Review and Meta-Analysis: Task-based fMRI Studies in Youths with Irritability"

Significant findings of task fMRI studies qualified for quantitative meta-analysis ($k$ = 28)

| Study | N | Significant Findings (Whole Brain and Region of Interest) | | | | | | | | Sex Differences | |
|---|---|---|---|---|---|---|---|---|---|---|---|
| | | | | | | MNI Coordinates | | | | | |
| | | Contrast | Cluster size | Cluster unit | Regions | x | y | z | High vs. low irritability | | Findings |
| Adleman et al. (2011)[42] | 82 | **WB: Acquisition vs. reversal phases (SMD vs. BD vs. HV)** | 1,674 | $mm^3$ | R superior temporal gyrus | 33 | 30 | -38 | ↑ | NA | |
| | | | 1,512 | | L superior temporal gyrus/insula | -49 | -14 | -19 | ↑ | | |
| | | | 945 | | R inferior temporal gyrus/middle frontal gyrus/fusiform | 66 | -16 | -27 | ↑ | | |
| | | | 810 | | L inferior temporal gyrus/middle frontal gyrus/fusiform | -51 | -6 | -39 | ↑ | | |
| | | | 783 | | R middle frontal gyrus | 34 | -1 | 57 | ↑ | | |
| | | | 729 | | R inferior temporal gyrus/middle frontal gyrus/fusiform | 49 | 0 | -37 | ↑ | | |
| | | WB: Incorrect vs. correct trials (SMD vs. BD vs. HV) | 2,484 | | Bilateral superior/medial frontal gyri | 2 | 9 | 73 | = | | |
| | | | 1,377 | | R inferior frontal gyrus/insula | 37 | 23 | 2 | ↓ | | |
| | | | 675 | | L fusiform/declive | -20 | -90 | -31 | ↑ | | |
| | | | 621 | | R cerebellum | 53 | -56 | -38 | = | | |
| | | WB: Acquisition vs. reversal phase x Incorrect vs. correct trials (SMD vs. BD vs. HV) | 1,512 | | R superior parietal lobule/precuneus | 28 | -79 | 55 | = | | |
| | | | 810 | | L superior temporal gyrus/insula | -52 | -11 | -13 | ↑ | | |
| | | | 675 | | L superior parietal lobule/precuneus | -25 | -72 | 62 | ↑ | | |
| | | | 675 | | R superior parental lobule/precuneus | 25 | -69 | 66 | ↑ | | |
| | | | 621 | | R caudate | 9 | 14 | -1 | ↑ | | |
| | | ROI: Incorrect vs. correct trials (SMD vs. BD vs. HV) | 621 | | R inferior frontal gyrus | 46 | 27 | 9 | ↓ | | |
| | | | 459 | | R caudate | 12 | 14 | -1 | ↓ | | |
| Aggensteiner et al. (2020)[38] | 177 | **WB: Negative faces vs. shapes (ODD and/or CD cases vs. HV)** | 59 | k | L frontal interior operculum | -51 | 11 | 23 | ↑ | NA^ | Not significant as a covariate. |
| | | | | | | -45 | 20 | 32 | ↑ | | |
| | | | 31 | | L superior occipital | -24 | -91 | 23 | ↑ | | |
| | | | 13 | | L inferior parietal gyrus | -36 | -61 | 53 | ↑ | | |
| | | | 10 | | L amygdala | -27 | -4 | -13 | ↑ | | |
| | | **WB: Positive faces vs. shapes (ODD and/or CD cases vs. HV)** | 45 | | R middle frontal | 33 | 26 | 50 | ↑ | | |
| | | | | | R superior frontal | 24 | 38 | 44 | ↑ | | |
| | | | | | | 24 | 23 | 62 | ↑ | | |
| | | | 32 | | R postcentral | 51 | -13 | -25 | ↑ | | |
| | | | | | | 54 | 2 | -28 | ↑ | | |

| | | | | | | | | | | |
|---|---|---|---|---|---|---|---|---|---|---|
| | | | 28 | | L occipital sup | -18 | -94 | 29 | ↑ | |
| | | | 26 | | R medial orbitofrontal | 6 | 50 | -7 | ↑ | |
| | | | | | | 15 | 41 | -10 | ↑ | |
| | | | | | | 6 | 38 | -13 | ↑ | |
| | | | 22 | | L olfactory | -3 | 11 | -10 | ↑ | |
| | | | | | | -6 | -1 | -13 | ↑ | |
| | | | 21 | | R inferior orbitofrontal | 24 | 14 | -25 | ↑ | |
| | | | | | | 18 | 17 | -19 | ↑ | |
| | | | 20 | | Not labelled | 9 | 2 | -10 | ↑ | |
| | | | 12 | | L medial oribitofrontal | -9 | 44 | -13 | ↑ | |
| | | ROI: Negative faces vs shapes (ODD and/or CD cases vs. HV) | 9 | | L amygdala | -27 | -4 | -13 | ↑ | |
| Bertocci et al. (2019)[43] | 96 | **WB: Happy vs. sad vs. angry faces (Irritability)** | -- | -- | R fusiform gyrus | 42 | -44 | -22 | ↑ | NA |
| Bubenzer-Busch et al. (2016)[18] | 34 (27) | **WB: Low aggression (Reactive aggression in HV)** | 1,365 | k | L middle temporal gyrus | -60 | -38 | 2 | ↑ | NA |
| | | | | | L middle temporal gyrus | -48 | -36 | -2 | ↑ | |
| | | | | | L superior temporal gyrus/temporal parietal junction | -46 | -44 | 26 | ↑ | |
| | | | | | L angular gyrus | -40 | -62 | 28 | ↑ | |
| | | | | | L middle temporal gyrus/temporal parietal junction | -50 | -52 | 26 | ↑ | |
| | | | | | L middle temporal gyrus/temporal parietal junction | -48 | -52 | 22 | ↑ | |
| | | | 640 | | R superior occipital gyrus | 28 | -68 | 32 | ↑ | |
| | | | | | R precuneus | 10 | -54 | 24 | ↑ | |
| | | | | | R cuneus | 14 | -62 | 26 | ↑ | |
| | | | | | R posterior cingulate cortex | 8 | -46 | 18 | ↑ | |
| | | | | | L precuneus | -6 | -60 | 14 | ↑ | |
| | | | | | L precuneus | -2 | -66 | 30 | ↑ | |
| | | | 319 | | R middle cingulate cortex | 2 | -40 | 38 | ↑ | |
| | | | | | L posterior cingulate cortex | -6 | -42 | 34 | ↑ | |
| | | | 143 | | R middle temporal gyrus/temporal parietal junction | 54 | -64 | 26 | ↑ | |
| | | | | | R middle temporal gyrus/temporal parietal junction | 56 | -58 | 22 | ↑ | |
| | | | | | R middle temporal gyrus/temporal parietal junction | 60 | -60 | 20 | ↑ | |
| Chaarani et al. (2020)#[19] | 320 | ROI: Successful vs. failed inhibition (Irritability) | -- | -- | R superior temporal gyrus | 30 | 17 | 8 | ↓ | NA |
| | | | | | L superior temporal gyrus | -61 | -13 | -1.5 | ↓ | |
| | | | | | R ventral precentral and postcentral gyri | 60 | 10 | 19.5 | ↓ | |
| Crum et al. (2020)[44] | 155 | **WB: Congruent vs. incongruent vs. view (Irritability)** | 116 | k | L posterior cingulate cortex | -5 | -30 | 35 | ↑ | No |
| | | | 84 | | R rostro-medial frontal gyrus | 5 | 55 | 20 | ↑ | |

| | | | | | | | | | | | |
|---|---|---|---|---|---|---|---|---|---|---|---|
| | | | 27 | | L anterior cingulate cortex | -4 | 29 | 25 | ↑ | | |
| | | WB: Congruent vs. incongruent vs. view (Irritability groups) | 292 | | L culmen extending to posterior cingulate cortex | -5 | -56 | -7 | ↑ | | |
| | | | 116 | | R superior frontal gyrus | 9 | 21 | 27 | ↑ | | |
| | | | 82 | | R inferior parietal lobule | 40 | -68 | 44 | ↑ | | |
| | | | 55 | | R rostro-medial frontal cortex | 9 | 53 | 20 | ↑ | | |
| | | | 27 | | L anterior cingulate cortex | -7 | 32 | 1 | ↑ | | |
| Deveney et al. (2013)[15] | 42 | **WB: Negative vs. positive feedback (SMD vs. HV)** | 296 | k | R posterior cingulate cortex | 17 | -41 | 12 | ↓ negative feedback | NA^ | Not significant as a covariate. |
| | | | 137 | | L posterior cingulate cortex | -20 | -58 | 33 | ↓ negative feedback | | |
| | | | 83 | | R postcentral gyrus and inferior parietal lobule | 63 | -27 | 36 | ↓ negative feedback | | |
| | | | 82 | | L supramarginal gyrus and inferior parietal lobule | -38 | -32 | 26 | ↓ negative feedback | | |
| | | | 74 | | L insula | -29 | -22 | 20 | ↓ negative feedback | | |
| | | | 65 | | R supramarginal gyrus | 46 | -55 | 29 | ↓ negative feedback | | |
| | | | 60 | | L insula | -42 | 9 | -1 | ↓ negative feedback | | |
| | | | 49 | | L parahippocampal gyrus | -14 | -11 | -20 | ↓ negative feedback | | |
| | | | 45 | | L postcentral/precentral gyrus | -42 | 16 | 23 | ↓ negative feedback | | |
| | | | 39 | | R precuneus | 14 | -61 | 41 | ↓ negative feedback | | |
| | | | 38 | | L cingulate/thalamus/caudate | -14 | -29 | 23 | ↓ negative feedback | | |
| Gatzke-Kopp et al. (2009)[17] | 30 | **WB: Noreward vs. reward (Externalizing vs. HV)** | 1,641 | $mm^3$ | L superior frontal gyrus | -16 | 54 | 24 | -- | NA | Boys only. |
| | | | | | R superior frontal gyrus | 16 | 34 | 32 | -- | | |
| | | | | | L middle frontal gyrus | -18 | 48 | 26 | -- | | |
| | | | | | R superior medial gyrus | 14 | 46 | 28 | -- | | |
| | | | 751 | | L angular gyrus | -46 | -66 | 28 | -- | | |
| | | | | | L middle temporal | -52 | -58 | 20 | -- | | |
| | | | | | L inferior parietal | -56 | -58 | 36 | -- | | |
| | | ROI: Nonreward vs. fixation (Externalizing vs. Controls) | 1,383 | | L caudate | -10 | 2 | 8 | ↑ | | |
| | | | | | L putamen | -20 | 20 | 2 | ↑ | | |
| | | | 1,342 | | R caudate | 30 | 16 | 2 | ↑ | | |
| | | | | | R putamen | 14 | 22 | 0 | ↑ | | |
| | | | 1,596 | | L caudate | -20 | -12 | 20 | ↑ | | |
| | | ROI: Reward vs. fixation (Externalizing) | | | L anterior cingulate cortex | -4 | 26 | -6 | ↓ nonreward | | |
| | | | | | R anterior cingulate cortex | 8 | 26 | -10 | ↓ nonreward | | |
| | | | 1,542 | | R caudate | 12 | 24 | 4 | ↑ | | |
| | | | | | R putamen | 26 | 16 | 2 | ↑ | | |
| Hodgdon et al. (2021)[45] | 31 | **WB: Reward received vs. reward blocked conditions** | 318 | k | R cuneus, R middle occipital gyrus, R lingual gyrus | 7 | -92 | 3 | ↑ | NA^ | Not significant as a covariate. |

| | | | | | | | | | | | |
|---|---|---|---|---|---|---|---|---|---|---|---|
| | | **(Irritability, child report)** | | | | | | | | | |
| | | | 125 | | Bilateral postieror cingulate cortex, bilateral cuneus, R precuneus | 3 | -72 | 12 | ↑ | | |
| | | | 100 | | R superior frontal gyrus, R medial frontal gyrus | 9 | -4 | 71 | ↑ | | |
| | | | 95 | | R temporal pole, R inferior frontal gyrus | 49 | 22 | -26 | ↑ | | |
| | | | 95 | | L lingual gyrus | -2 | -94 | -15 | ↑ | | |
| | | **WB: Reward received vs. reward blocked during n+1 anticipation period (Irritability, child report)** | 71 | | R lingual gyrus, R inferior occipital gyrus | 22 | -93 | -11 | ↑ after reward blocked; ↓ after reward received | | |
| Ibrahim et al. (2019)[46] | 57 | **WB: Fear vs. calm faces (ASD + DB vs. HV)** | 1,488 | $mm^3$ | L superior temporal gyrus | -64 | -54 | 18 | ↑ (ASD + DB) | NA | |
| | | | | | L middle temporal gyrus | -54 | -16 | -20 | ↑ (ASD + DB) | | |
| | | | | | L inferior temporal gyrus | -50 | -10 | -30 | ↑ (ASD + DB) | | |
| | | | 711 | | L precentral gyrus | -42 | -20 | 64 | ↑ (ASD + DB) | | |
| Karim et al. (2017)#[26] | 51 | WB: Negative vs. neutral clips (Irritability x Age interaction) | 245 | | L superior medial frontal | -8 | 56 | 30 | ↑ | NA | |
| | | | 239 | | L cerebellum declive | -10 | -64 | -26 | ↑ | | |
| | | | 222 | | L superior frontal gyrus | -14 | 38 | 42 | ↑ | | |
| | | | 197 | | | -8 | -64 | -28 | ↑ | | |
| | | | 176 | | L caudate | -12 | 20 | 14 | ↑ | | |
| | | | 133 | | L middle frontal gyrus | -26 | 32 | 42 | ↑ | | |
| | | | 119 | | L brainstem (medulla) | -4 | -44 | -46 | ↑ | | |
| | | | 116 | | | -22 | -64 | -36 | ↑ | | |
| | | | 115 | | | -26 | -70 | -38 | ↑ | | |
| | | | 95 | | L thalamus | -12 | -12 | 6 | ↑ | | |
| | | | 71 | | L putamen | -14 | 16 | -2 | ↑ | | |
| | | | 68 | | R brainstem (medulla) | 2 | -46 | -48 | ↑ | | |
| | | | 68 | | L cerebellum pyramis | -12 | -70 | -40 | ↑ | | |
| Kircanski et al. (2018)[47] | 197 | **WB: Threat congruent vs. threat incongruent trials (Irritability)** | 9,125 | $mm^3$ | L inferior parietal lobule/L postcentral gyrus/L superior parietal lobule | -46 | -28 | 47 | ↑ | NA^ | No significant effect of sex was reported as a covariate. |
| | | | 4,625 | | L caudate/left lentiform nucleus | -10 | 8 | 10 | ↑ | | |
| | | | 2,438 | | L dorsolateral prefrontal cortex | -27 | 39 | 34 | ↑ | | |
| | | | 2,172 | | L inferior parietal lobe | 53 | -62 | 38 | ↑ | | |
| | | | 1,563 | | L insula | 38 | -2 | -7 | ↑ | | |
| | | | 1,563 | | R dorsolateral prefrontal cortex | 21 | 40 | 37 | ↑ | | |
| | | | 1,453 | | R caudate/R lentiform nucleus | 17 | 12 | -6 | ↑ | | |
| | | | 1,359 | | L ventrolateral prefrontal cortex | -34 | 54 | 6 | ↑ | | |
| | | WB: Threat vs. neutral trials (Negative affectivity) | 1,234 | | R dorsomedial nucleus of thalamus/ L dorsomedial nucleus of thalamus | 5 | -10 | 7 | ↑ | | |

| | | | | | | | | | | | |
|---|---|---|---|---|---|---|---|---|---|---|---|
| | | ROI: Threat congruent vs. threat incongruent trials (Irritability) | 1,063 | | L amygdala | -21 | -1 | -29 | ↑ | | |
| Kryza-Lacombe et al. (2020a)[11] | 45 | WB: Angry-neutral vs. happy-neutral condition vs. sad-neutral vs. neutral-neutral face-emotion pairs (Irritability) | 8,045 | k | L postcentral gyrus, R precuneus | -56 | -32 | 47 | -- | NA^ | Not significant as a covariate. |
| | | | 1,455 | | L middle occipital gyrus, L cuneus | -10 | -92 | -17 | -- | | |
| | | | 1,364 | | R middle frontal gyrus, R superior frontal gyrus | 42 | 26 | 35 | -- | | |
| | | | 1,080 | | R superior temporal gyrus | 43 | -6 | -33 | -- | | |
| | | | 386 | | R cerebellar tonsil | 32 | -22 | -48 | -- | | |
| | | | 383 | | R pyramis | 8 | -71 | -38 | -- | | |
| | | | 347 | | L parahippocampal gyrus, L middle temporal gyrus | -37 | -15 | -21 | -- | | |
| | | | 286 | | L cingulate gyrus | -18 | -8 | 29 | -- | | |
| | | | 271 | | R lingual gyrus | 24 | -94 | -27 | -- | | |
| | | | 215 | | R lentiform nucleus | 43 | -39 | -3 | -- | | |
| | | | 202 | | R declive, R tuber | 54 | -54 | -25 | -- | | |
| | | | 155 | | L cingulate gyrus | -10 | -49 | 24 | -- | | |
| | | | 139 | | L declive | -53 | -53 | -28 | -- | | |
| | | | 139 | | R inferior frontal gyrus | 45 | 33 | 5 | -- | | |
| | | | 135 | | L transverse temporal gyrus | -36 | -31 | 10 | -- | | |
| | | | 108 | | L uncus, L inferior frontal gyrus | -25 | 25 | -28 | -- | | |
| | | | 103 | | R pyramis, R cerebellar tonsil | 49 | -58 | -44 | -- | | |
| | | | 97 | | L pyramis | -8 | -84 | -41 | -- | | |
| | | | 91 | | R superior temporal gyrus, R insula | 45 | -24 | -2 | -- | | |
| | | | 78 | | L medial frontal gyrus, L superior frontal gyrus | -19 | 42 | 19 | -- | | |
| | | | 76 | | R cuneus | 17 | -92 | 31 | -- | | |
| | | | 74 | | R inferior frontal gyrus | 52 | 42 | 4 | -- | | |
| | | | 72 | | L precentral gyrus | -61 | -8 | 40 | -- | | |
| | | | 63 | | R inferior parietal lobule, R postcentral gyrus | 37 | -32 | 45 | -- | | |
| | | **WB: Angry-neutral vs. happy-neutral vs. sad-neutral vs. neutral-neutral face-emotion pairs x Congruent vs. incongruent trials (Irritability)** | 3,724 | | Bilateral postcentral gyri, L precentral gyri, L superior parietal lobule | 27 | -31 | 78 | ↑ happy-incongruent trials; ↓ angry faces | | |
| | | | 553 | | R pyramis | 18 | -69 | -40 | ↑ happy-incongruent trials; ↓ angry faces | | |
| | | | 382 | | R inferior parietal lobe | 48 | -47 | 57 | ↑ happy-incongruent trials; ↓ angry faces | | |
| | | | 150 | | L lingual gyrus | -26 | -71 | -26 | ↑ happy-incongruent trials; ↓ angry faces | | |
| | | | 144 | | R cingulate gyrus | 9 | -27 | 38 | ↑ happy-incongruent trials; ↓ angry faces | | |
| | | | 110 | | R superior frontal gyrus | 24 | 63 | 3 | ↑ happy-incongruent trials; ↓ angry faces | | |

| | | | | | | |
|---|---|---|---|---|---|---|
| | 110 | R postcentral gyrus, R inferior parietal lobule | 53 | -32 | 47 | ↑ happy-incongruent trials; ↓ angry faces |
| | 105 | R middle temporal gyrus | 54 | -25 | -22 | ↑ happy-incongruent trials; ↓ angry faces |
| | 105 | R precuneus, R inferior parietal lobule | 28 | -80 | 46 | ↑ happy-incongruent trials; ↓ angry faces |
| | 101 | R superior temporal gyrus | 47 | -39 | 2 | ↑ happy-incongruent trials; ↓ angry faces |
| | 94 | R postcentral gyrus | 41 | -28 | 40 | ↑ happy-incongruent trials; ↓ angry faces |
| | 92 | R parahippocampal gyrus | 43 | 10 | -21 | ↑ happy-incongruent trials; ↓ angry faces |
| | 81 | L inferior occipital gyrus | -25 | -99 | -6 | ↑ happy-incongruent trials; ↓ angry faces |
| | 78 | R superior frontal gyrus | 20 | 64 | 27 | ↑ happy-incongruent trials; ↓ angry faces |
| | 70 | R inferior frontal gyrus | 45 | 31 | 8 | ↑ happy-incongruent trials; ↓ angry faces |
| WB: Congruent vs. incongruent trials (Irritability) | 1,203 | L postcentral gyrus, precentral gyrus | -47 | -37 | 66 | -- |
| | 99 | L precuneus, L cingulate gyrus | -14 | -45 | 24 | -- |
| WB: Angry-neutral vs. happy-neutral vs. sad-neutral vs. neutral-neutral face-emotion paris (Irritability x Anxiety) | 1,883 | Bilateral postcentral gyri, L superior/inferior parietal lobule | -4 | -50 | 77 | -- |
| WB: (Irritability) | 955 | L cerebellar tonsil | 10 | -82 | -41 | -- |
| | 757 | L postcentral gyrus | -19 | -58 | 74 | -- |
| | 109 | R precuneus | 7 | -58 | 41 | -- |
| | 83 | R postcentral gyrus | 10 | -60 | 82 | -- |
| | 82 | R postcentral gyrus, middle temporal gyrus | -51 | -42 | -19 | -- |
| WB: Congruent vs. incongruent trials (Irritability x Anxiety) | 723 | R pyramis, R declive, R uvula | 22 | -68 | -42 | -- |
| WB: (Irritability x Anxiety) | 321 | L precuneus, L superior parietal lobule | -19 | -70 | 60 | -- |
| | 214 | R precuneus, R superior parietal lobule | 14 | -66 | 61 | -- |
| | 141 | L posterior cingulate | -18 | -63 | 9 | -- |
| | 134 | L cuneus | -5 | -86 | 29 | -- |
| | 117 | R postcentral gyrus | 62 | -26 | 45 | -- |
| | 113 | L cuneus | -15 | -92 | 31 | -- |
| | 87 | R inferior parietal lobule | 35 | -48 | 71 | -- |
| | 67 | L fusiform gyrus | -40 | -39 | -8 | -- |
| | 67 | R postcentral gyrus | 44 | -24 | 66 | -- |
| | 60 | R lingual gyrus, R cuneus | 24 | -95 | -9 | -- |
| WB: Angry-neutral vs. happy-neutral vs. sad-neutral vs. neutral-neutral face-emotion pairs x Congruent vs. incongruent trials (Irritability x Anxiety) | 233 | L superior parietal lobule | -35 | -62 | 64 | ↑ (low and high anxiety) |

| | | | | | | | | | | | |
|---|---|---|---|---|---|---|---|---|---|---|---|
| | | | 65 | | R cuneus | 21 | -96 | 30 | ↑ (low and high anxiety) | | |
| Kryza-Lacombe et al. (2020b)[48] | 120 | **WB: Happy vs. fearful vs. sad vs. neutral faces (Irritability)** | 98 | k | L middle frontal gyrus | -36 | 32 | 35 | ↓ fearful, happy | NA^ | Not significant as a covariate. |
| | | | 38 | | L inferior frontal gyrus | -46 | 11 | 7 | ↓ happy | | |
| | | WB: (Irritability main effect) | 36 | | L precuneus | -9 | -66 | 42 | ↑ | | |
| Kryza-Lacombe et al. (2021)[37] | 52 | WB: Reward anticipation period (Irritability main effect) | 157 | k | R cuneus | 15 | -78 | 9 | ↑ | No | |
| | | | 129 | | R uncus | 34 | -3 | -50 | ↓ | | |
| | | | 127 | | R cerebellar tonsil | 13 | -53 | -55 | -- | | |
| | | | 107 | | R striatum | 26 | 10 | -5 | ↑ | | |
| | | | 74 | | L uncus | -23 | -4 | -43 | ↓ | | |
| | | **WB: Reward vs. no-reward conditions during reward anticipation period (Irritability)** | 91 | | L inferior parietal lobule | -49 | -39 | 59 | ↓ nonreward | | |
| | | | 77 | | L cuneus | -10 | -105 | -6 | ↑ nonreward | | |
| | | WB: Performance feedback period (Irritability main effect) | 152 | | R cerebellar tonsil | 13 | -53 | -53 | -- | | |
| | | | 86 | | L caudate | -12 | -1 | 25 | ↑ | | |
| | | | 66 | | L postcentral gyrus | -60 | -15 | 15 | ↑ | | |
| | | **WB: Reward vs. no-reward conditions during performance feedback (Irritability)** | 355 | | L precuneus and cuneus | -3 | -82 | 49 | ↑ | | |
| | | | 288 | | R lingual gyrus | 9 | -77 | -20 | ↑ | | |
| | | | 189 | | R fusiform gyrus | 43 | -57 | -33 | ↑ | | |
| | | | 179 | | L cuneus | -3 | -80 | 5 | ↑ | | |
| | | | 117 | | Bilateral culmen | -3 | -43 | -2 | ↑ | | |
| | | | 114 | | R temporal parietal junction | 50 | -40 | 19 | ↑ | | |
| | | WB: Reward/hit vs. reward/miss vs. no reward/hit vs. no reward/miss (Irritability) | 202 | | R precuneus | 30 | -73 | 44 | ↑ | | |
| | | | 100 | | Bilateral cingulate gyri | -4 | -41 | 22 | ↑ | | |
| | | | 89 | | R middle and inferior occipital gyrus | 34 | -82 | 7 | ↑ | | |
| | | WB: Reward vs. no-reward conditions x Reward/hit vs. reward/miss vs. no reward/hit vs. no reward/miss (Irritability) | 296 | | L temporal parietal junction | -48 | -35 | 7 | ↑ | | |
| | | | 173 | | R precuneus | 15 | -71 | 51 | ↑ | | |
| | | | 130 | | L fusiform gyrus | -61 | -45 | -26 | ↑ | | |
| | | | 117 | | L striatum | -22 | -7 | -4 | ↑ | | |
| | | | 96 | | L inferior parietal lobule | -35 | -44 | 45 | ↑ | | |
| | | | 74 | | L postcentral gyrus | -56 | -19 | 50 | ↑ | | |
| | | | 70 | | R pyramind | 26 | -81 | -40 | ↑ | | |

| | | | | | | | | | | | |
|---|---|---|---|---|---|---|---|---|---|---|---|
| Liuzzi et al. (2020)[20] | 19 | **WB: Congruent vs. incongruent condition (Irritability)** | 74 | k | L amygdala/L parahippocampal gyrus, L uncus | -32 | -9 | -28 | ↓ congruent | NA^ | Gender was analysed as a covariate as an additional analysis. The main effect of irritability in the bilateral cingulate gyri, L posterior cingulate, and in the R thalamus became not significant. |
| | | WB: (Irritability) | 471 | | L middle frontal gyrus, L superior frontal gyrus, L inferior frontal gyrus | -38 | 38 | 5 | ↑ | | |
| | | | 179 | | Bilateral postcentral gyri, L precentral gyrus, L superior parietal lobule | -9 | 46 | -4 | ↑ | | |
| | | | 139 | | R lentiform nucleus/striatum | 16 | 7 | 8 | ↑ | | |
| | | | 105 | | Bilateral declive, R uvula | 14 | -70 | -29 | ↑ | | |
| | | | 68 | | L declive, L tuber | -44 | -68 | -28 | ↑ | | |
| Pagliaccio et al. (2017)[49] | 83 | **WB: Time point interactions (Diagnosis) - average BOLD** | 162 | k | R postcentral gyrus | 25 | -37 | 57 | ↑ | No | |
| | | | 85 | | R declive | 36 | -75 | -31 | = | | |
| | | | 73 | | L postcentral gyrus | -30 | -36 | 57 | ↑ | | |
| | | | 72 | | R medial frontal gyrus | 9 | -23 | 57 | ↑ | | |
| | | | 48 | | L thalamus | -3 | -21 | -1 | ↑ | | |
| | | | 39 | | R parahippocampal gyrus | 31 | -35 | -15 | ↓ | | |
| | | | 39 | | L posterior cingulate | -6 | -56 | 24 | = | | |
| | | | 29 | | R cerebellar tonsil | 24 | -44 | -58 | ↑ | | |
| | | | 25 | | R superior frontal gyrus | 2 | 10 | 54 | = | | |
| | | WB: Time point interactions (Diagnosis) - reaction time BOLD | 64 | | R paracentral lobule | 2 | -40 | 66 | ↑ before stimulus; = peak | | |
| | | | 43 | | L percuneus | -23 | -57 | 46 | = before stimulus; ↓ peak | | |
| | | | 41 | | R superior parietal lobule | 11 | -69 | 63 | ↑ before stimulus; ↓ peak | | |
| | | | 27 | | R fusiform gyrus | 53 | -39 | -19 | ↑ before stimulus; ↓ peak | | |
| | | | 25 | | R culmen | 53 | -40 | -42 | ↑ before stimulus; = peak | | |
| | | WB: Congruent vs. incongruent conditions x Timepoint interactions (Diagnosis) - reaction time BOLD | 72 | | L precuneus | -12 | -65 | 58 | ↑ before stimulus; ↓ peak during congruent trials | | |
| | | | 40 | | L supramarginal gyrus | -55 | -42 | 36 | ↑ before stimulus; ↓ peak during congruent trials | | |
| | | | 7 | | L precuneus | -12 | -72 | 46 | ↑ before stimulus; ↓ peak during | | |

| | | | | | | | | | | | |
|---|---|---|---|---|---|---|---|---|---|---|---|
| | | | | | | | | | congruent trials | | |
| | | | 6 | | R middle frontal gyrus | 18 | -13 | 65 | ↑ before stimulus; ↓ peak during congruent trials | | |
| | | | 19 | | R superior occipical gyrus | 36 | -79 | 28 | ↑ before stimulus; ↓ peak during congruent trials | | |
| | | | 9 | | L inferior frontal gyrus | -46 | 3 | 29 | ↑ before stimulus; ↓ peak during congruent trials | | |
| Perlman et al. (2015)[50] | 54 | WB: (Group) | 1,750 | $mm^3$ | R postcentral gyrus | 57 | -23 | 41 | ↑ | No | |
| | | **WB: Win vs. loss (Irritability vs. HV)** | 907 | | R parahippocampal gyrus/posterior cingulate | 11 | -48 | 2 | ↓ winning; ↑ losing | | |
| | | | 901 | | R anterior cingulate cortex | 12 | 36 | 19 | ↑ winning; ↓ losing | | |
| | | | 643 | | L middle frontal gyrus | -1 | 58 | 25 | ↑ winning; ↓ losing | | |
| | | | 578 | | R superior frontal gryus | 12 | 51 | 27 | ↑ | | |
| Stoddard et al. (2017)[51] | 115 | **WB: Happy vs. angry vs. fearful face response (Irritability)** | 91 | k | L postcentral gyrus | -25 | -35 | 58 | ↑ | NA | |
| | | | 86 | | R fusiform gyrus | 43 | -54 | -21 | ↑ | | |
| | | | 63 | | R middle occipital gyrus | 35 | -84 | 8 | ↑ | | |
| | | | 55 | | L pulvinar | -23 | -29 | 9 | ↑ | | |
| | | | 54 | | R pulvinar | 25 | -25 | 5 | ↑ | | |
| | | | 48 | | R midcingulate | 13 | -29 | 41 | ↑ | | |
| | | | 43 | | R superior occipital | 32 | -79 | 25 | ↑ | | |
| Strenziok et al. (2011)[52] | | **WB: Aggressive vs. non-aggressive conditions (Trait anger)** | -- | -- | L ventromedial prefrontal cortex | -3 | 50 | -7 | ↓ | NA | |
| Thomas et al. (2012)%[27] | 57 | WB: Neutral vs. angry faces (Group main effect) | 454 | k | L posterior cingulate | -6 | -61 | 9 | ↓ (BPD, SMD) | No | A significant main effect of sex in the L amygdala ROI (smaller average beta weights for males than females); a main effect of sex at a trend level in the L middle/superior frontal gyrus (smaller average beta weights for females than males); no significant group x sex effect. |
| | | WB: Neutral vs. happy faces (Group main effect) | 407 | | R inferior parietal lobule | 51 | -61 | 43 | ↑ (SMD) | | |
| | | | 368 | | L middle occipital gyrus, fusiform gyrus | -39 | -62 | -6 | ↑ (SMD) | | |
| | | | 341 | | R middle occipital gyrus, cuneus | 34 | -90 | 6 | ↑ (SMD) | | |
| | | | 254 | | L middle superior frontal gyrus | -33 | -15 | 52 | ↑ (SMD) | | |

| | | | | | | | | | | | |
|---|---|---|---|---|---|---|---|---|---|---|---|
| Thomas et al. (2013)[36] | 53 | **WB: Angry vs. fearful vs. neutral expressions (Diagnosis)** | 149 | k | R anterior cingulate gyrus | -5 | 11 | 34 | ↑ | NA | |
| | | | 92 | | R posterior insula | 37 | -16 | 9 | ↑ | | |
| | | | 69 | | L inferior parietal lobe | -31 | -55 | 37 | ↑ | | |
| | | | 65 | | R posterior cingulate gyrus | -5 | -30 | 37 | ↑ | | |
| | | | 45 | | L posterior insula | -39 | -22 | 13 | ↑ | | |
| | | | 30 | | L posterior cingulate gyrus | 2 | -37 | 44 | ↑ | | |
| | | | 27 | | L anterior cingulate gyrus | 5 | 11 | 34 | ↑ | | |
| Thomas et al. (2014)[53] | 60 | WB: Nonaware vs. aware conditions (Diagnosis) | 94 | k | R middle occipital gyrus | 25 | -85 | -3 | ↑ | NA | |
| | | | 26 | | L middle occipital gyrus | -24 | -93 | -3 | ↑ | | |
| | | | 22 | | L middle occipital gyrus | -36 | -68 | -3 | ↑ | | |
| | | **WB: Anger vs. fear vs. happy vs. neutral vs. no face (Diagnosis)** | 84 | | L precentral gyrus | -60 | -16 | 30 | = | | |
| | | | 52 | | R posterior cingulate | 16 | -34 | 20 | ↑ angry (SMD) | | |
| | | | 31 | | R superior temporal gyrus | 44 | -7 | 8 | ↑ angry (SMD, BD); ↑ happy (SMD) | | |
| | | | 24 | | R middle occipital gyrus | 34 | -53 | 4 | ↑ angry (SMD) | | |
| | | | 23 | | L medial frontal gyrus | -10 | 50 | 16 | = | | |
| Tseng et al. (2016)[12] | 37 | **WB: Angry vs. happy vs. neutral vs. noface (SMD vs. HV)** | 187 | k | L insula | -32 | -16 | 19 | ↓ happy | NA^ | Not significant as a covariate. |
| | | | 87 | | L culmen | -26 | -35 | -29 | ↑ angry; ↓ happy | | |
| | | | 60 | | L culmen/parahippocampal gyrus | -28 | -43 | -18 | ↑ angry; ↓ happy | | |
| | | | 59 | | R parahippocampal gyrus | 36 | -17 | -27 | ↑ angry; ↓ happy | | |
| | | | 52 | | L declive | -20 | -77 | -28 | ↑ angry; ↓ happy | | |
| | | | 42 | | R thalamus | 17 | -13 | 1 | ↓ happy | | |
| | | | 37 | | L cerebellar lingual | -5 | -44 | -18 | ↑ angry; ↓ happy | | |
| | | | 34 | | R superior temporal gyrus | 65 | -20 | -1 | ↑ angry | | |
| | | | 33 | | R culmen | 10 | -25 | -30 | ↑ angry; ↓ happy | | |
| | | | 26 | | R cingulate gyrus | 14 | -36 | 30 | ↑ angry; ↓ happy | | |
| | | WB: Aware vs. non-aware (SMD vs. HV) | 65 | | L ventromedial prefrontal cortex | -4 | 41 | -10 | ↑ | | |
| Tseng et al. (2019)[16] | 195 | **WB: After rigged vs. positive feedback (Irritability)** | 13,594 | $mm^3$ | Bilateral cingulate gyrus, R superior frontal gyrus | 9 | 17 | 44 | ↑ | No | Gender did not moderate most of the main results except in the inferior parietal lobule, pre- and post-central gyri, and insula. Irritability was associated with increased activation in younger boys, and decreased activation in older boys. |
| | | | 5,844 | | R middle frontal gyrus | 36 | 14 | 43 | ↑ | | |
| | | | 2,469 | | L middle frontal gyrus | -32 | 21 | 36 | ↑ | | |
| | | | 2,422 | | R caudate, thalamus | 11 | -19 | 18 | ↑ | | |

| | | | | | | | | | | |
|---|---|---|---|---|---|---|---|---|---|---|
| | | | 1,422 | | R dorsolateral prefrontal cortex | 47 | 30 | 26 | ↑ | |
| | | | 1,250 | | R cuneus | 9 | -79 | 17 | ↑ | |
| | | | 1,094 | | R precuneus | 18 | -73 | 45 | ↑ | |
| | | | 1,047 | | L middle frontal gyrus | -34 | -4 | 50 | ↑ | |
| | | | 594 | | R inferior frontal gyrus | 56 | 28 | 3 | ↑ | |
| | | | 531 | | L precentral and postcentral gyri | -40 | -21 | 42 | ↑ | |
| | | | 469 | | L parahippocampal gyrus | -17 | -39 | -10 | ↑ | |
| | | | 453 | | L caudate | -10 | 7 | 22 | ↑ | |
| | | | 406 | | R superior temporal gyrus | 41 | -51 | 17 | ↑ | |
| | | | 344 | | R precentral gyrus | 60 | 9 | 2 | ↑ | |
| | | | 266 | | L precentral gyrus | -64 | 0 | 13 | ↑ | |
| | | | 203 | | L cingulate gyrus | -7 | -18 | 44 | ↑ | |
| | | | 203 | | L superior frontal gyrus | -7 | 7 | 66 | ↑ | |
| | | ROI: After rigged vs. positive feedback (Irritability) | -- | | Bilateral caudate | 14 | 13 | 11 | ↑ | |
| | | | -- | | L putamen | -26 | 3 | -1 | ↑ | |
| Tseng et al. (2021)[54] | 59 | **WB: Explicit memory vs. threat appraisal (Irritability, parent report)** | 1225 | k | L cerebellum | -22 | -79 | -36 | ↓ | NA |
| | | | 854 | | L pre-/post-central gyrus | -59 | -19 | 15 | ↓ | |
| | | | 277 | | L ventrolateral prefrontal cortex | -37 | 43 | -7 | ↓ | |
| | | | 227 | | L superior temporal gyrus | -52 | -46 | 5 | ↓ | |
| | | | 153 | | R superior temporal gyrus | 60 | -4 | 0 | ↓ | |
| | | | 147 | | R precuneus | 9 | -53 | 66 | ↓ | |
| | | | 99 | | L parahippocampal gyrus | -28 | -23 | -27 | ↓ | |
| | | | 96 | | R primary motor cortex | 51 | -14 | 47 | ↓ | |
| | | | 95 | | R inferior parietal lobule | 38 | -62 | 36 | ↓ | |
| | | | 84 | | R posterior insula | 50 | -35 | 18 | ↓ | |
| | | | 77 | | R amygdala | 20 | -7 | -12 | ↓ | |
| | | | 73 | | R inferior parietal lobule | 50 | -57 | 41 | ↓ | |
| | | | 72 | | R middle temporal gyrus | 49 | -60 | -1 | ↓ | |
| | | | 64 | | L primary motor cortex | -3 | -28 | 60 | ↓ | |
| | | | 58 | | R dorsolateral prefrontal cortex | 29 | 40 | 42 | ↓ | |
| | | | 50 | | R cerebellum | 45 | -61 | -47 | ↓ | |
| | | | 48 | | L cuneus | -4 | -85 | 21 | ↓ | |
| | | | 41 | | L lingual gyrus | -5 | -97 | -18 | ↓ | |
| | | | 40 | | R cuneus | 11 | -84 | 39 | ↓ | |
| | | | 34 | | L middle temporal gyrus | -53 | 12 | -31 | ↓ | |
| | | | 30 | | R superior parietal lobule | 11 | -74 | 63 | ↓ | |
| | | | 29 | | L inferior parietal lobule | -29 | -52 | 50 | ↓ | |
| | | WB: Explicit memory vs. threat appraisal (Irritability, child report) | 170 | | L postcentral gyrus | -63 | -20 | 26 | ↓ | |
| | | | 148 | | R cerebellum | 20 | -74 | -34 | ↓ | |
| | | | 123 | | L parahippocampal gyrus | -28 | -23 | -27 | ↓ | |
| | | | 111 | | L inferior parietal lobule | -36 | -59 | 41 | ↓ | |
| | | | 105 | | R fusiform gyrus | 29 | -61 | -24 | ↓ | |
| | | | 98 | | R inferior parietal lobule | 38 | -59 | 35 | ↓ | |
| | | | 88 | | R pre-/post-central gyrus | 57 | -16 | 41 | ↓ | |

| | | | | | | | | | | |
|---|---|---|---|---|---|---|---|---|---|---|
| | | | 87 | | L ventrolateral prefrontal cortex | -58 | 18 | 13 | ↓ | |
| | | | 78 | | L ventrolateral prefrontal cortex | -42 | 43 | -15 | ↓ | |
| | | | 61 | | L premotor cortex | -62 | 1 | 7 | ↓ | |
| | | | 61 | | L cerebellum | -20 | -82 | -36 | ↓ | |
| | | | 50 | | R middle frontal gyrus | 32 | 6 | 32 | ↓ | |
| | | | 46 | | L middle temporal gyrus | -53 | 11 | -28 | ↓ | |
| | | | 44 | | R amygdala | 25 | 2 | -27 | ↓ | |
| | | | 44 | | L temporoparietal junction | -54 | -51 | 4 | ↓ | |
| | | | 40 | | L midcingulate cortex | -5 | -16 | 44 | ↓ | |
| | | | 33 | | R fusiform gyrus/cerebellum | 32 | -38 | -30 | ↓ | |
| | | | 30 | | L superior temporal gyrus | -52 | -4 | -6 | ↓ | |
| | | | 29 | | L V2 | -11 | -94 | 21 | ↓ | |
| Wiggins et al. (2016)[13] | 71 | WB: Fearful vs. happy vs. angry emotion x 0% vs. 50% vs. 75% vs. 100% intensity (Diagnosis, Irritability) - modelled cubically | 300 | k | R posterior superior temporal sulcus/temporal parietal junction | 34 | -48 | 24 | ↑ middle intensity, fearful and angry faces (DMDD) | NA |
| | | | 128 | | L temporal parietal occipital junction | -47 | -65 | 7 | ↑ middle intensity, fearful and angry faces (DMDD) | |
| | | | 84 | | R temporal pole | 32 | 25 | -28 | ↑ middle intensity, fearful and angry faces (DMDD) | |
| | | | 75 | | L superior temporal sulcus | -64 | -26 | 9 | ↑ middle intensity, fearful and angry faces (DMDD) | |
| | | | 63 | | L amygdala | -19 | -2 | -17 | ↑ middle intensity faces | |
| | | | 47 | | L lingual gyrus | -20 | -55 | 2 | ↑ middle intensity, fearful and angry faces (DMDD) | |
| | | | 45 | | R lateral prefrontal cortex | 43 | 28 | 6 | ↑ middle intensity faces | |
| | | | 40 | | R postcentral gyrus | 32 | -49 | 66 | ↑ middle intensity faces | |
| | | WB: Fearful vs. happy vs. angry emotion x 0% vs. 50% vs. 75% vs. 100% intensity (Diagnosis, Irritability) - modelled quadratically | 2,760 | | L precuneus, cingulate, lingual gyrus | -5 | -40 | 68 | -- | |
| | | | 790 | | L posterior superior temporal sulcus, supramarginal gyrus | -47 | -71 | -11 | -- | |

| | | | | | | | | | | |
|---|---|---|---|---|---|---|---|---|---|---|
| | | | 663 | | R superior temporal gyrus/rolandic operculum | 67 | -27 | 17 | -- | |
| | | | 190 | | R cuneus, calcarine gyrus | 21 | -72 | 24 | -- | |
| | | | 182 | | R caudate nucleus, putamen | 7 | 22 | 5 | -- | |
| | | | 178 | | L thalamus | -3 | -16 | 13 | -- | |
| | | | 174 | | R cerebellum | 21 | -46 | -63 | -- | |
| | | | 170 | | L amygdala/hippocampus | -26 | 5 | -17 | -- | |
| | | | 131 | | R fusiform, inferior temporal gyrus | 37 | -53 | -24 | -- | |
| | | | 130 | | L postcentral gyrus | -52 | -16 | 47 | -- | |
| | | | 96 | | L middle temporal gyrus | -32 | -63 | 12 | -- | |
| | | | 81 | | R temporal occipital junction | 45 | -67 | 8 | -- | |
| | | | 74 | | R cerebellum | 45 | -51 | -50 | -- | |
| | | | 67 | | L middle/superior occipital gyrus | -19 | -83 | 20 | -- | |
| | | | 66 | | L calcarine gyrus, cuneus | -11 | -75 | 24 | -- | |
| | | | 62 | | R precentral gyrus | 31 | -21 | 38 | -- | |
| | | | 53 | | L postcentral gyrus | -25 | -28 | 42 | -- | |
| | | | 52 | | L cerebellar vermis | -2 | -66 | -41 | -- | |
| | | | 45 | | L cerebellum | -17 | -72 | -24 | -- | |
| | | | 41 | | L superior orbital gyrus | -9 | 25 | -20 | -- | |
| | | **WB: Fearful vs. happy vs. angry emotion x 0% vs. 50% vs. 75% vs. 100% intensity (Diagnosis, Irritability) - modelled linearly** | 54 | | R temporal pole | 32 | 25 | -26 | -- | |
| | | | 53 | | R middle orbital gyrus | 2 | 64 | 9 | -- | |
| | | | 46 | | R medial prefrontal cortex | 15 | 52 | 16 | -- | |
| Yang et al. (2017)[55] | 48 (31) | WB: Biological vs. scrambled motion (HV vs ASD with low ODD) | 4.35 | z | R middle occipital gyrus | 44 | -88 | 6 | ↓ | NA |
| | | | 3.97 | | R middle temporal gyrus | 56 | -72 | 0 | ↓ | |
| | | | 3.68 | | R inferior temporal gyrus | 58 | -64 | -14 | ↓ | |
| | | | 3.61 | | R inferior occipital gyrus | 46 | -86 | -2 | ↓ | |
| | | | 2.56 | | R fusiform gyrus | 42 | -54 | -16 | ↓ | |
| | | WB: Biological vs. scrambled motion (HV vs. ASD with high ODD) | 4.13 | | R angular gyrus | 34 | -58 | 42 | ↓ | |
| | | | 3.38 | | R inferior parietal gyrus | 36 | -54 | 42 | ↓ | |
| | | | 3.37 | | R middle temporal gyrus | 44 | -68 | 16 | ↓ | |
| | | | 3.15 | | R inferior temporal gyrus | 42 | -58 | -8 | ↓ | |
| | | | 3.06 | | R supramarginal gyrus | 38 | -42 | 42 | ↓ | |
| | | | 2.82 | | R fusiform gyrus | 34 | -44 | -4 | ↓ | |
| | | | 2.82 | | R middle occipital gyrus | 34 | -62 | 38 | ↓ | |
| | | **WB: Fixation vs. biological motion (ASD with high vs. low ODD)** | 3.46 | | L angular gyrus | -42 | -58 | 40 | ↓ | |
| | | | 3.14 | | L anterior cingulate, paracingulate gyrus | -2 | 42 | 6 | ↓ | |
| | | | 2.96 | | L superior frontal gyrus, medial | -8 | 62 | 8 | ↓ | |
| | | | 2.85 | | R anterior cingulate, paracingulate gyrus | 6 | 36 | 8 | ↓ | |

| | | | | | |
|---|---|---|---|---|---|
| 2.81 | L inferior parietal gyrus | -50 | -58 | 46 | ↓ |
| 2.69 | L supramarginal gyrus | -56 | -52 | 32 | ↓ |

*Notes.* # Studies qualified for the quantitative analysis but excluded from the main analysis because significant results were found only in the ROI analysis,[19] and no significant clusters were reported for any task interaction effects with irritability (without age interaction).[26] % [27] was excluded from any GingerALE analyses as only significant main effects of irritability were found upon data extraction. ^ Studies did not test for sex moderation or interaction effects but analyzed as a covariate. Adjusted sample sizes for studies that conducted the contrast analysis in specific subgroups of interest were presented in parentheses. Boldfaced contrasts were included in the main analysis in GingerALE. ASD = Autism Spectrum Disorder; BD = Bipolar Disorder; CD = Conduct Disorder; DBD = Disruptive Behavior Disorder; DMDD = Disruptive Mood Dysregulated Disorder; fMRI = functional magnetic resonance imaging; HV = healthy volunteers; MNI = Montreal Neurological Institute; ODD = Oppositional Defiant Disorder; SMD = Severe Mood Dysregulation; L/R = Left/Right hemisphere; WB = whole-brain analysis; ROI = region-of-interest analysis; -- = Not reported.
